# Targeting androgen regulation of TMPRSS2 and ACE2 as a therapeutic strategy to combat COVID-19

**DOI:** 10.1101/2020.10.16.342782

**Authors:** Qu Deng, Reyaz ur Rasool, Ronnie M. Russell, Ramakrishnan Natesan, Irfan A. Asangani

## Abstract

Epidemiological data showing increased severity and mortality of COVID-19 in men suggests a potential role for androgen in SARS-CoV-2 infection. Here, we present evidence for the transcriptional regulation of SARS-CoV-2 host cell receptor ACE2 and TMPRSS2 by androgen in mouse and human cells. Additionally, we demonstrate the endogenous interaction between TMPRSS2 and ACE2 in human cells and validate ACE2 as a TMPRSS2 substrate. Further, Camostat – a TMPRSS2 inhibitor, blocked the cleavage of pseudotype SARS-CoV-2 surface Spike without disrupting TMPRSS2-ACE2 interaction. Thus providing evidence for the first time a direct role of TMPRSS2 in priming the SARS-CoV-2 Spike, required for viral fusion to the host cell. Importantly, androgen-deprivation, anti-androgens, or Camostat attenuated the SARS-CoV-2 S-mediated cellular entry. Together, our data provide a strong rationale for clinical evaluations of TMPRSS2 inhibitors, androgen-deprivation therapy/androgen receptor antagonists alone or in combination with antiviral drugs as early as clinically possible to prevent COVID-19 progression.

## Introduction

The ongoing COVID-19 (Coronavirus disease 2019) pandemic caused by Severe Acute Respiratory Syndrome Coronavirus 2 (SARS-CoV-2) infection is global health crisis (Morens and Fauci, 2020; Wang et al., 2020). As of January 2021, over 100 million cases of COVID-19 and more than two million deaths have been recorded (https://coronavirus.jhu.edu). Epidemiological data have shown that males are disproportionally affected, being slightly more likely to be infected than females and accounting for most severely ill cases and higher fatality. This is compounded by older age and comorbidities such as diabetes, cardiovascular diseases, obesity, hypertension and cancer (Williamson et al., 2020). This sex disparity with respect to increased morbidity and mortality in men suggests a potential role for the male hormone androgen in SARS-CoV-2 infection and host response (Stopsack et al., 2020; Wadman, 2020). The host immune response to SARS-CoV-2 potentiates a hyper-inflammatory cytokine storm involving the upregulation of tumor necrosis factor-α (TNF-α), interleukin 1β (IL-1 β), interleukin-6 (IL-6), monocyte chemoattractant protein-1 (MCP-1), and macrophage inflammatory protein 1-α (MIP1α), which is associated with COVID-19 severity and mortality(Mehta et al., 2020; Moore and June, 2020; Vabret et al., 2020; Zhang et al., 2020). Tumor cell-intrinsic and microenvironment associated immune cell response to inflammation promote the development and progression of several types of solid cancer, including prostate cancer (de Bono et al., 2020). Therefore, it is reasonable to hypothesize that the inflammatory signaling accompanying severe COVID-19 disease could cause cancer progression. Therefore, early intervention of COVID-19 management should be considered in cancer patients who are more susceptible to SARS-CoV-2 infection due to their immunosuppressive state.

SARS-CoV-2 entry into the host cell is mediated by trimers of the transmembrane spike (S) glycoprotein. The S trimer binds to the cellular receptor angiotensin converting enzyme 2 (ACE2) on the surface of target cells and mediates subsequent viral uptake and fusion dependent on processing by host proteases (Hoffmann et al., 2020c). Several host proteases, including TMPRSS2, cathepsin B and L, and furin have been suggested to promote the entry of SARS-CoV-2; how they process the Spike protein remains to be determined (Coutard et al., 2020; Shang et al., 2020; Walls et al., 2020). In relevant target cells, the cleavage of S by TMPRSS2 activates the protein for membrane fusion via extensive, irreversible conformational changes (Millet and Whittaker, 2015; Walls et al., 2020; Walls et al., 2017). As a result, SARS-CoV-2 entry into susceptible cells is a complex process that requires the concerted action of ACE2 receptor-binding and TMPRSS2 proteolytic processing of the S protein to promote virus-cell fusion. Despite their well-documented role in SARS-CoV-2 entry, the nature of endogenous ACE2 and TMRPSS2 interactions in host cells is lacking.

TMRPSS2 is a widely studied androgen-regulated gene associated with prostate cancer (Afar et al., 2001). It contributes to prostate cancer pathogenesis by way of aberrantly driving oncogenic transcription. More than half of all prostate cancers in men of European ancestry harbor a gene fusion that juxtaposes the androgen receptor (AR) regulatory promoter/enhancer elements of *Tmprss2* in front of the ETS family transcription factors, most commonly *Erg* (Tomlins et al., 2005). Although ERG is not normally regulated by androgen, this somatic gene rearrangement results in oncogenic ERG expression under androgen receptor signaling. Further, TMRPSS2 is known to promote metastatic spread of prostate cancer through HGF activation (Lucas et al., 2014). ACE2 is a zinc-dependent metalloprotease and is expressed at high levels in the heart, testis, and kidney and at lower levels in various tissue (Gheblawi et al., 2020). Therefore, identifying molecular mechanisms governing the expression of TMPRSS2 and ACE2 could potentially explain the observed disparity in SARS-CoV-2 infection-associated morbidity and mortality in men.

In this study, we provide evidence for the AR-mediated transcriptional regulation of TMPRSS2 and ACE2 in mice and human prostate and lung cells. Specifically, androgen deprivation in mice by castration, or anti-androgen treatment *in vitro* led to a reduction in TMPRSS2 and ACE2 transcript and protein levels. Furthermore, TMPRSS2 and ACE2 were found to interact in *cis* in prostate and lung cells, and the inhibition of TMPRSS2 protease activity by Camostat blocked Spike priming. Finally, androgen-deprivation, anti-androgens, or Camostat treatment attenuated the SARS-CoV-2 pseudotype entry in lung and prostate cells, suggesting that these drug combinations could be effective in attenuating COVID-19 disease progression in men with or without prostate cancer.

## Results

### Systemic androgen deprivation affects TMPRSS2 and ACE2 expression in mice

The poor clinical outcome of COVID-19 in men suggests a potential underlying androgen-related cause. The role of male sex hormone androgen in regulating the SARS-CoV-2 host cell receptor ACE2 and co-receptor TMPRSS2 has been speculated (La Vignera et al., 2020; Stopsack et al., 2020; Wadman, 2020). To evaluate the effect of androgen deprivation on TMPRSS2 and ACE2 expression in major tissues that are the primary sites for SARS-CoV-2 infection(Ziegler et al., 2020), we performed surgical castration in adult male mice (n=3). One week post-castration, we harvested the lung, small intestine, heart and kidney for qRT-PCR, immunoblotting, and immunohistochemistry analysis. Seminal vesicles and prostate were used as positive controls for systemic androgen deprivation. Tissues from the corresponding mock castrated male (n=3) and female (n=3) served as controls (Fig. 1A). First, we determined the expression of *Tmprss2, Ace2* and *Ar* in the control tissues. Each was expressed at varying degrees, with *Ar* and *Tmprss2* being the highest in seminal vesicles, and *ACE2* in the small intestine (Supplementary Fig. 1A). Interestingly, a survey of adult human male tissue mRNA expression via bulk RNA-seq from the GTEx consortium(Consortium et al., 2017) demonstrated a similar expression pattern in prostate and small intestine for *AR, TMPRSS2*, and *ACE2* (Supplementary Fig. 1B), suggesting a potentially identical mechanism of regulation in man and mice. Next, as expected, the *Tmprss2* expression was highly androgen-responsive in the seminal vesicles of the castrated males compared to the control males. (Fig. 1B). Except for the small intestine and the lung which showed a slight downward trend, levels of *Tmprss2* in the heart (with very low basal expression), and the kidney did not display any change upon castration. However, *Ace2* expression displayed a significant downregulation in the lung and the small intestine, which was similar to the levels found in female mice, and upregulation in the kidney tissues in the castrated male compared to controls. Remarkably, *Ace2* expression was also reduced in the hormone-sensitive seminal vesicles upon castration (Fig. 1B), suggesting a potential role for androgen in regulating *Ace2* expression in these tissues. In concordance with the transcripts, the immunoblot analysis using total protein extracts from tissues demonstrated a reduced TMRPSS2 and ACE2 protein levels in the lung and the small intestine. The reduction in protein levels was much more dramatic than the transcript levels. In contrast, the kidney tissue displayed an increase in the ACE2 protein, but not TMPRSS2, corroborating the transcript data (Fig. 1C). Next, we performed immunohistochemistry to identify the specific cell types in these tissues that display altered expression for TMPRSS2 and ACE2 proteins upon androgen deprivation. In these experiments, DU145 and VCaP prostate cancer xenograft tissues served as a negative control for AR, TMPRSS2, and ACE2 respectively, which showed a clear lack of staining (Supplementary Fig. 2A). As expected, based on the transcript data, reduced staining for TMPRSS2, ACE2, and AR protein in hormone-sensitive seminal vesicles was observed in the castrated group (Supplementary Fig. 2B). Further, the hormone-responsive prostate luminal epithelial cells showed reduced staining for AR and TMPRSS2 in response to castration (Fig. 1D). We also observed a small minority population of epithelial cells <5% staining for ACE2 on the apical surface of the prostate epithelium, which was absent in the castrated group. The identical minimal co-expression pattern of ACE2 and TMPRSS2 was also identified in human prostate tissues, where club cells that constitute a mere 0.07% of total prostate epithelial cells and resemble club cells in the lung were double-positive(Henry et al., 2018; Montoro et al., 2018; Song et al., 2020). We observed no morphological changes in response to castration in lung, small intestine and kidney tissue through H&E staining. Similar to transcripts, the AR, ACE2, and TMPRSS2 proteins were detected in all of the tested tissue types at varying degrees (Fig. 1E). A strong staining for TMPRSS2 was detected exclusively in the lung respiratory bronchiole epithelium, with minimal staining of type II alveolar cells, which is consistent with the expression pattern observed in human tissue (Stopsack et al., 2020). The staining intensity, especially in type II alveolar cells, was reduced in the castrated group. The ACE2 staining could be observed in the lung parenchyma, both type I and type II alveolar cells and the bronchiole epithelium apical surface of the castrated males showed relatively lower expression (Fig. 1E and Supplementary Fig. 2C). Next, the small intestine mucosa from the control group showed positive cytoplasmic and nuclear AR staining, which were reduced in the castrated group. TMPRSS2 displayed high staining in the crypt and lower portion of the villi with the exception in the goblet cells, and the staining gradually diminished on the top of the villi. ACE2 was mainly present on the apical surface of the enterocytes on the top part of the villi. These expression patterns mirror data from human small intestinal tissue, especially ileum and jejunum (Hamming et al., 2004; Ziegler et al., 2020). The abundance of double-positive cells was largely reduced in the castrated male tissue due to the reduction in both ACE2 and TMPRSS2 levels. Concerning the kidney, the AR staining was detected in the tubular cells, which was reduced upon castration. TMPRSS2 showed positivity mainly in the proximal tubules compared to other cells, which was reduced in the castrated group (Muus et al., 2020). Finally, a homogenous ACE2 expression was detected across kidney tissue on the apical surface of the tubular cells. In line with the kidney’s role in the renin-angiotensin-aldosterone system, the ACE2 intensity was strongly elevated in the castrated males responding to reduced blood pressure brought out by androgen deprivation (Dubey et al., 2002). Together, these analyses clearly demonstrate that the androgen deprivation has an effect on the expression of TMPRSS2 and ACE2, particularly in the lung, small intestine, and kidney. Though there was a discrepancy concerning transcript and protein levels of TMPRSS2 in the tested tissues - particularly in the kidney, post-transcriptional regulation of TMPRSS2 by androgen regulated microRNA could not be ruled out (Nersisyan et al., 2020).

**Figure 1:**
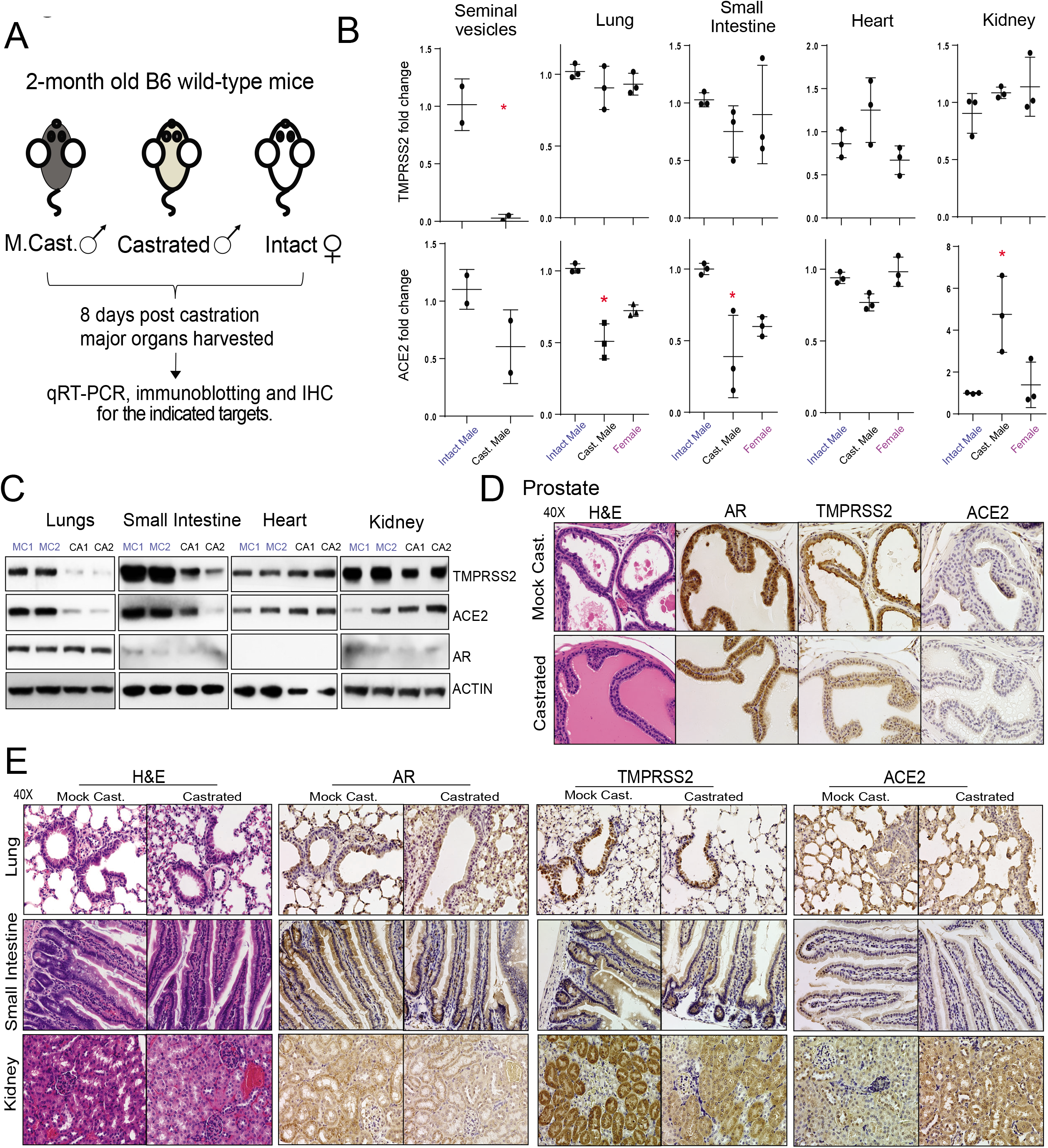
Effect of androgen deprivation by castration on the expression of TMPRSS2 and ACE2 in adult male mice. **(A)** Schematic depicting the castration experiment in 8-9 week-old wild-type C57BL/6J mice. **(B)** Varying effect of systemic androgen deprivation on the transcription of TMPRSS2 and ACE2 in major organs. qRT-PCR analysis for TMPRSS2 and ACE2 transcripts in the indicated organs from mock, castrated male and normal females. Highly hormone-responsive seminal vesicles from the male mice served as a positive control for the effect of androgen deprivation on the target genes. **(C)** Immunoblot analysis showing the levels of TMPRSS2, ACE2, and AR proteins in the indicated organs from two separate mock vs. castrated males. β-Actin served as a loading control. **(D)** Immunohistochemistry analysis of the indicated target protein in the prostate from mock and castrated males. Note the reduced TMPRSS2 staining in the castrated group, and lack/sparse ACE2 staining in both groups. **(E)** As in d for the indicated organs. Staining for AR was observed in all the tested organs. Note the reduced TMPRSS2 staining in the bronchial epithelium of lung, columnar epithelium of small intestine, and proximal convoluted tubules in the kidney, in the castrated group. Also, note the reduced ACE2 staining in alveolar epithelium of the lungs and columnar epithelium of small intestine, and increased staining in the kidney from the castrated group. Also, see Supplementary Figure 1-2. * p < 0.05 (Student’s t-test).

### AR regulates TMPRSS2 and ACE2 expression in human prostate and lung cells

The observation that systemic androgen deprivation affects the expression of TMPRSS2 and ACE2 in hormone-sensitive tissues such as prostate and seminal vesicles, but also in major organs such as the lungs, small intestine, and kidney led us to investigate the direct role of AR in the transcriptional regulation of these two critical genes beyond the prostate. Although the upstream enhancer of *Tmprss2* is a well-known target for AR (Asangani et al., 2014), we sought to identify whether regulatory elements of *Ace2* are also occupied by AR. We analyzed publicly available AR ChIP-seq data for AR-binding sites within 50 kb of the transcription start sites (TSSs) of the *Tmprss2* and *Ace2* genes in mouse prostate, comparing castration and castration followed by testosterone injection for 3 days (Pihlajamaa et al., 2014). As expected, multiple AR peaks were observed in the upstream of the *Tmprss22* TSS in the testosterone-treated prostate, which was also present in DHT (dihydrotestosterone) treated VCaP and LNCaP human prostate cell lines (Fig. 2a). Interestingly, the upstream region of *ACE2* also showed AR binding in mouse as well as in human prostate cells (Fig. 2A and Supplementary Fig. 3A). Motif analysis of the AR occupied regions revealed the presence of androgen response elements (ARE), as well as binding sites for other steroid receptors such as GR (Glucocorticoid Receptor) and PR (Progesterone Receptor), and pioneering factor such as FOXA1 (Supplementary Fig. 3A), suggesting potential regulation of ACE2 by other steroid hormones. Next, we wanted to investigate whether TMPRSS2 and ACE2 expression in human cells responds to androgen deprivation or anti-androgen treatment. We cultured LNCaP cells in steroid-deprived conditions for 2, 4, and 6 days and assessed the expression of TMPRSS2 and ACE2 compared to cells grown in steroid-proficient condition. As one of the most androgen-responsive gene, TMPRSS2 transcript and protein levels showed downregulation in the steroid-deprived cells compared to their steroid-proficient controls (Fig. 2B). This was accompanied by a concomitant decrease in the levels of phosphoSer81-AR (pAR), an active chromatin-bound form of AR (Rasool et al., 2019). More importantly, the ACE2 transcript and the protein levels were also downregulated in the steroid-deprived cells. Treatment with the second-generation anti-androgen enzalutamide or AR degrader ARD-69 (Han et al., 2019) led to a time-dependent reduction in the TMPRSS2 and ACE2 expression that was accompanied by reduced pAR levels in LNCaP cells (Fig. 2C). Since lung is one of the primary site of SARS-CoV-2 infection causing severe disease, we tested whether lung cells respond to androgen-deprivation and anti-androgen treatment with respect to the expression of TMPRSS2 and ACE2. First, we screened a panel of lung cancer cell lines and found H460 cells positive for AR, TMPRSS2 and ACE2 (Fig. 2D). Similar to that observed in LNCaP cells, H460 displayed a marked downregulation of TMPRSS2 and ACE2 accompanied by reduced pAR levels upon androgen-deprivation, or AR targeted therapy (Fig. 2E). In a complementary experiment, stimulation with DHT in LNCaP and H460 cells led to increased pAR and concomitant increase in TMPRSS2 and ACE2 protein levels (Supplementary Fig. 3B). Interestingly, androgen induced AR dependent upregulation of TMPRSS2 has been demonstrated in other lung cancer cell lines (Mikkonen et al., 2010). These data clearly demonstrate a potential role of AR in regulating SARS-CoV-2 receptor and co-receptor in prostate and lung cells.

**Figure 2:**
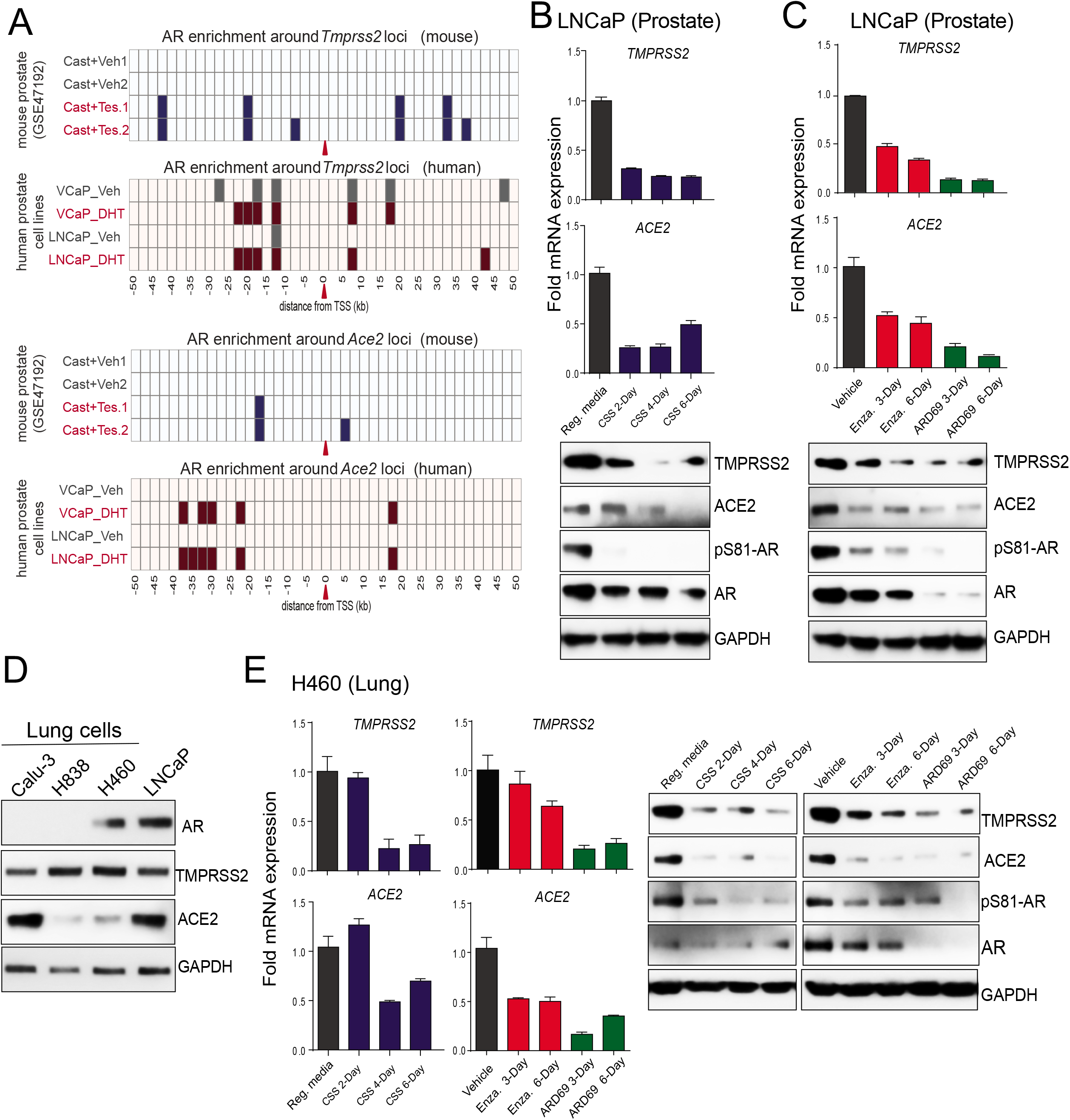
AR-mediated transcriptional regulation of TMPRSS2 and ACE2 in human prostate and lung cells. **(A)** *TMPRSS2*, and *ACE2* loci display enhanced AR-binding upon testosterone stimulation. Shown are AR ChIP-seq peaks around *TMPRSS2* and *ACE2* locus (the red arrow indicates the TSS) in the vehicle and testosterone-treated conditions in mouse prostate tissue (GSE47192) and human prostate cell lines. **(B)** Androgen deprivation, anti-androgen or AR-degradation results in the loss of TMPRSS2 and ACE2 expression. Top, qRT-PCR analysis for TMPRSS2 and ACE2 transcripts in human prostate cells – LNCaP, grown in the indicated conditions; Reg. media – Regular media with 10% serum, CSS – charcoal striped serum, Enza. – Enzalutamide at 25 μM, ARD69 – AR degrader at 250nM. Bottom, immunoblot analysis for the indicated protein. GAPDH was used the loading control. **(C)** Immunoblot showing the expression of AR, TMPRSS2, ACE2 and GAPDH protein in a panel of human lung cell lines, LNCaP was used as the positive control. **(D)** qRT-PCR, and immune blot analysis as in b for human lung cell line H460. Also, see Supplementary Figure 3.

### TMPRSS2 interacts with ACE2 in prostate and lung cells

Having identified the androgen dependency of TMPRSS2 and ACE2 expressions *in vivo* in mice and *in vitro* in human cell lines, we turned our attention to their function concerning SARS-CoV-2 Spike (SARS-2-S) priming. Though TMPRSS2 is implicated in priming of SARS-2-S protein (Hoffmann et al., 2020c), it is unclear whether this occurs in association with ACE2. Further, TMPRSS2-mediated enhancement of SARS virus entry has been shown to accompany ACE2 cleavage (Shulla et al., 2011). Therefore, we studied whether TMPRSS2 and ACE2 physically associate in an endogenous setting and examined the effect of TMPRSS2 protease inhibition on their interaction and cleavage of ACE2. TMPRSS2 is composed of a cytoplasmic region (aa 1-84), a transmembrane region (aa 85-105), and an extracellular region (aa 106-492). The latter is composed of three domains: the LDLRA (LDL receptor class A) domain (residues 112-149) – that forms a binding site for calcium, the SRCR (scavenger receptor cysteine-rich) domain (aa 149-242) - involved in the binding to other cell surface or extracellular proteins, and the Peptidase S1 domain (aa 256-489), which contains the protease active site: residues H296, D345, and S441 (Fig. 4A). An auto-cleavage site at aa 255-256 has been shown to allow shedding of the extracellular region of TMPRSS2 (Afar et al., 2001). The ACE2 protein is 805 aa long and is composed of a short cytoplasmic region, a transmembrane region, and an extracellular N-terminal region that contain zinc metallopeptidase domain (Fig. 3A). TMPRSS2 mediated cleavage of ACE2 at its N-terminus (residues 697 to 716) was shown to be required for ACE2 to interact with SARS-S protein in 293T overexpression system (Heurich et al., 2014; Shulla et al., 2011). Our experiments show that overexpression of TMPRSS2 in HEK293-T cells that stably express ACE2 led to ACE2 cleavage, resulting in reduced levels of the ∼120 kDa full-length form and increased levels of a ∼20 kDa C-terminal fragment (Fig. 3B), as previously documented (Shulla et al., 2011). Importantly, the treatment of cells with Camostat – a TMPRSS2 serine protease inhibitor (Kawase et al., 2012) (Fig. 3C), entirely blocked ACE2 cleavage, suggesting ACE2 as a *bona fide* substrate of TMPRSS2 (Shulla et al., 2011). Intriguingly, Camostat treatment did not result in any change in ACE2 full-length levels in LNCaP cells (Supplementary Fig. 4A). This observation led us to address the existence of TMPRSS2-ACE2 complex in the endogenous setting. Towards that, we determined the physical interaction between TMPRSS2 and ACE2 protein using N-and C-terminal specific antibodies (Fig. 3A). Reciprocal immunoprecipitation experiments showed an endogenous association between TMPRSS2 and ACE2 in LNCaP prostate cells (Fig. 3D). Likewise, Calu-3 lung cells demonstrated a physical interaction between TMPRSS2 and ACE2 (Fig. 3E). Along with the full-length 50 kDa TMPRSS2, a smaller approximately 38 kDa variant was found to interact with ACE2 prominently in both the cell lines. Notably, the treatment of cells with Camostat did not reduce the amount of ACE2 being pulled down by TMPRSS2 and vice-versa, suggesting the endogenous TMPRSS2-ACE2 complex to be stable and is not dependent on proteolytic cleavage. A similar endogenous ACE2-TMPRSS2 interaction was observed in AR-positive H460 lung cells (Supplementary Fig. 4B). These data demonstrate for the first time an endogenous association between TMPRSS2 and ACE2 in prostate and lung cells.

**Figure 3:**
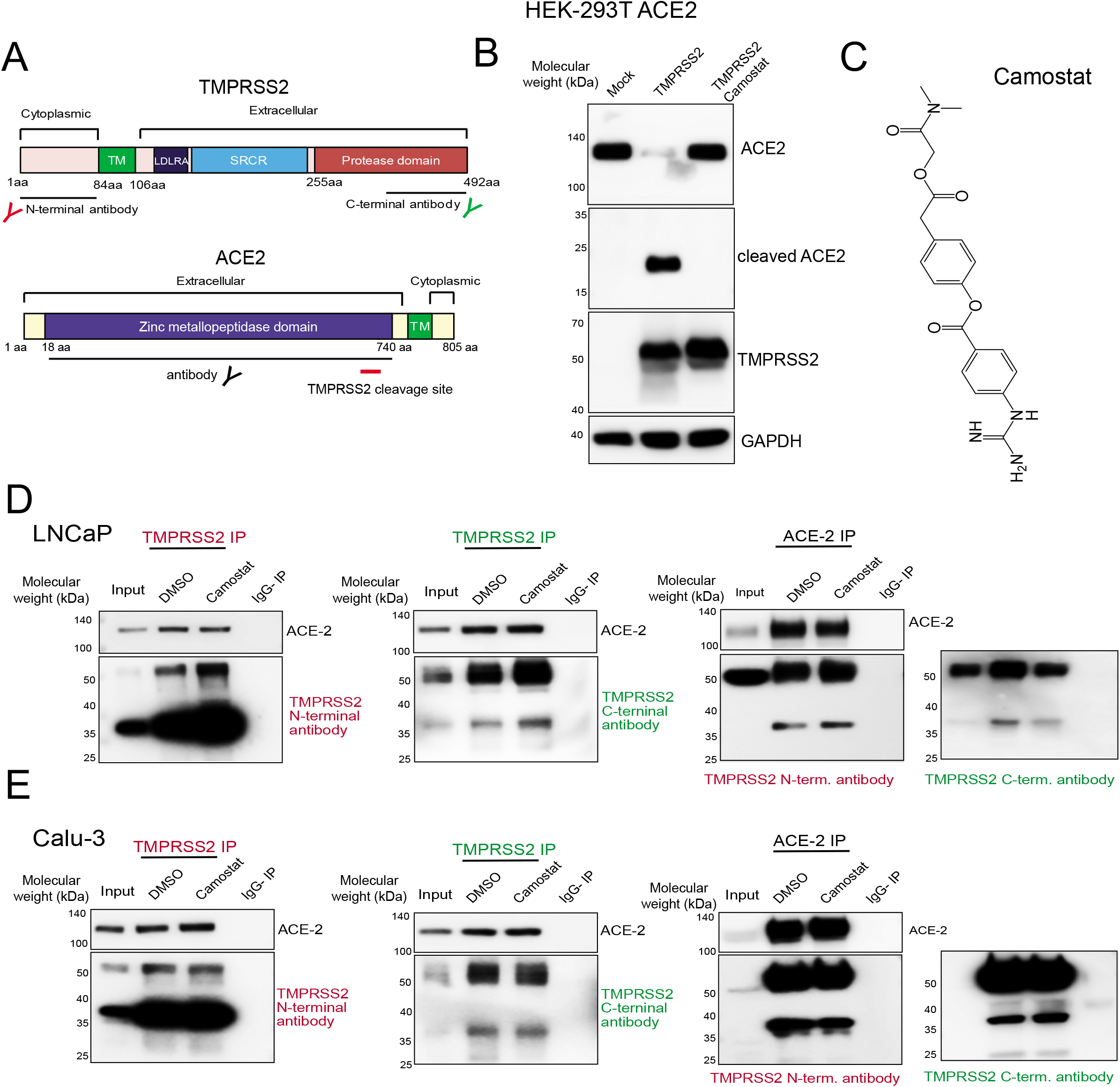
TMPRSS2 physically interacts with ACE2 in prostate and lung cells. **(A)** Schematic showing the domain structure of human TMPRSS2 and ACE2 protein. The TMRPSS2 cleavage site on ACE2 is shown with red bar. Epitopes recognized by the antibodies used in the immunoprecipitation are indicated. **(B)** TMPRSS2 cleaves ACE2. HEK 293T ACE2 cells were transfected with TMPRSS2 encoding plasmid, and 6h post transfection the cells were treated with DMSO or Camostat (250 μM). 48h post-transfection, proteins were extracted and used for immunoblotting with anti-ACE2, anti-TMPRSS2, and anti-GAPDH antibody. **(C)** Chemical structure of Camostat. **(D)** TMPRSS2 and ACE2 physically interacts and is not disrupted by Camostat – a TMPRSS2 inhibitor. Reciprocal immunoprecipitation using the indicated antibodies with LNCaP protein extracts. Cells were treated with DMSO or Camostat (250 μM) for 24hr followed by protein extraction. Note, along with the full-length ∼50kDa TMPRSS2 protein, the presence of a novel approximately 38kDa cleaved form detected in both N-terminal and C-terminal TMPRSS2 antibody pulldown and ACE2 pulldown. IgG pulldown served as a negative control; inputs were 5-10%. **(E)** As in D with Calu-3 protein extracts. Also, see Supplementary Figure 4.

**Figure 4:**
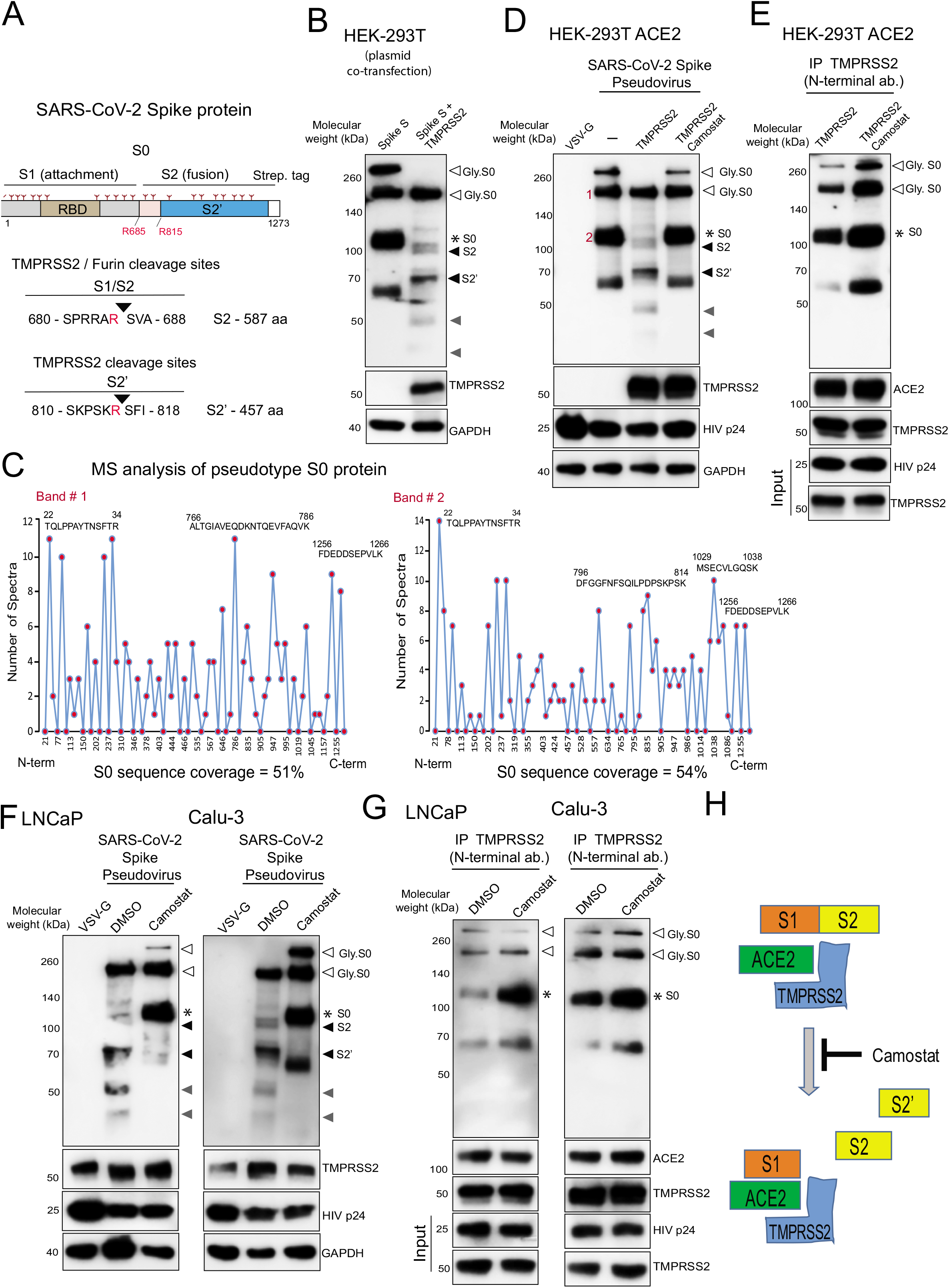
TMPRSS2 in complex with ACE2 cleaves SARS-CoV-2 Spike protein. **(A)** Schematic representation of the expression construct of full-length SARS-CoV-2 spike (S0) protein that has S1 and S2 segments involved in virus attachment and fusion, respectively. Segments of S1 include RBD, receptor-binding domain; S1/S2 cleavage site; S2 has an S2’ cleavage site. Tree-like symbol denotes glycans, and the C terminus contains 2x Strep-tag. The TMPRSS2 cleavage site on S1/S2 and S2’ site is indicated with the resulting length of the S2 and S2’ protein. Cleaved S2 and S2’ fragment lengths are shown. **(B)** TMPRSS2 cleaves glycosylated SARS-2-S protein. HEK 293T cells were transfected with SARS-2-S plasmid with or without TMPRSS2 encoding plasmid. 48h post-transfection, proteins were extracted and used for immunoblotting with anti-Streptavidin, anti-TMPRSS2, and anti-GAPDH antibody. The open arrow and * indicates the glycosylated non-glycosylated uncleaved S0 protein, respectively. The filled arrow black indicates TMPRS2 cleaved S2 and S2’. The filled gray arrow indicates TMPRSS2 cleaved potential S2 and S2’ fragments arising from non-glycosylated S0. **(C)** MS analysis of pseudotype incorporated S0 protein. SARS-2-S pseudovirus was directly subjected to streptavidin-IP, followed by MS analysis for sequencing. Histograms show peptide coverage for the two prominent bands detected with in IP-western (See supplementary figure 4). **(D)** Pseudotype SARS-CoV-2 Spike is cleaved by TMPRSS2, which is blocked by Camostat. HEK 293T ACE2 overexpressing cells were transfected with TMRPSS2 plasmid, 48h later cells were pretreated with 500 μM Camostat for 4h followed by pseudovirus inoculation by spinoculation for 1h. Next, 4h post-spinoculation recovery total lysates were prepared and used for immunoblotting with the indicated antibody. VSV-G containing pseudotype virus served as a negative control, and HIVp24 served as a positive control for viral inoculation. **(E)** Camostat blocks TMRPSS2 mediated cleavage of S0 without disrupting the interaction between TMPRSS2-ACE2-Spike S complex. Lysate from **c** was used to immunoprecipitate TMPRSS2 using N-terminal antibody followed by immunoblotting for the indicated target. Note the increased pulldown of uncleaved Spike S0 by the TMPRSS2 and the complete absence of S2, S2’, and other cleaved fragments in both DMSO and Camostat lane. **(F)** Camostat reverse the endogenous TMPRSS2 mediated Spike cleavage in human prostate and lung cells. The LNCaP and Calu-3 cells were pretreated with Camostat (500 μM) for 4hr followed by pseudotype VSV-G or SARS-CoV-2 particle spinoculation for 1h and processed as in **D. (G)** Camostat blocks TMRPSS2 mediated cleavage of S0 and S2 without disrupting the interaction between TMPRSS2-ACE2-Spike S complex in LNCaP and Calu-3 cells. Lysate from **e** was used to immunoprecipitate TMPRSS2 using N-terminal antibody followed by immunoblotting for the indicated target. Note the increased pulldown of Spike S0 by the TMPRSS2 and the lack of S2, S2’ and other fragments in the Camostat lane. **(H)** Schematic showing the interaction between Spike S, ACE2 and TMPRSS2 leading to cleavage of Spike S into S1, S2 and S2’ segments. S0/S1 continues to bind tightly through its RBD to ACE2 that is in complex with TMPRSS2, whereas upon cleavage the S2 and S2’ gets dissociated from the TMPRSS2-ACE2 complex. Also, see Supplementary Figure 5.

### TMPRSS2 inhibition blocks S priming without affecting its interaction with ACE2

Next, we addressed the role of TMPRSS2 on priming of SARS-CoV-2 Spike (SARS-2-S) – a heavily glycosylated transmembrane protein (Cai et al., 2020; Ke et al., 2020) (Fig. 4A). The unprocessed Spike S0 protein encoded by the SARS-CoV-2 genome is 1,273 amino acid long. It is incorporated into the viral membrane and facilitates viral entry into target cells. Similar to SARS CoV and MERS CoV Spike proteins, SARS-CoV-2 Spike consists of a proprotein convertase (PPC) motif at the S1/S2 boundary cleaved by furin protease and TMPRSS2 into two fragments – the receptor-binding fragment S1 and the membrane fusion fragment S2 (Hoffmann et al., 2020b; Hoffmann et al., 2020c; Iwata-Yoshikawa et al., 2019; Shang et al., 2020; Shulla et al., 2011; Zhou et al., 2015). Though the role of TMPRSS2 in SARS-2-S priming has been suggested (Hoffmann et al., 2020c), direct evidence demonstrating the cleaving of SARS-2-S by endogenous TMPRSS2 is lacking. Therefore, we performed experiments to specifically address the role of ectopically expressed or endogenous TMPRSS2 in SARS-2-S activation. Transfection of codon optimized plasmid encoding SARS-2-S protein and 2xStrep.tag (Gordon et al., 2020) in HEK293-T showed a distinct ∼180 kDa, and a ∼280 kDa band which is likely the differentially glycosylated S0 protein. The two other bands at ∼120 kDa and a ∼60 kDa could be the non-glycosylated fragments (Fig. 4B). The S1 fragment could not be detected in the immunoblots as the streptavidin-tag is located at the C-terminal end of the constructs. The co-transfection with TMPRSS2 plasmid led to cleavage of S0 into distinct bands corresponding to S2, and S2’ (Fig. 4B). Interestingly, the ∼280 kDa band and ∼120 kDa was fully processed by TMPRSS2, suggesting the S1/S2 site on glycosylated and non-glycosylated S0 protein could be equally targeted by TMPRSS2 for cleavage. The smaller fragments observed is most likely the TMPRSS2 cleaved S2 and S2’ from the non-glycosylated S0. To further demonstrate the action of TMPRSS2 on S activation, we performed spinoculation with pseudotype virus that contain replication deficient human immunodeficiency virus (HIV) particles bearing SARS-2-S proteins in TMPRSS2 negative HEK293-T ACE2 cells. First, the Mass spectrometry-based sequencing of pseudotype incorporated SARS-2-S identified the prominent ∼180 kDa and ∼120 kDa fragments as full-length S0 proteins, suggesting the presence of uncleaved Spike on the pseudotype (Fig. 4C and Supplementary Fig. 5A-B). Additionally, we tested and found that HEK293-T ACE2 cells have superior uptake of HIV pseudotype SARS-2-S compared to HEK293-T cells, confirming ACE2 as the major receptor SARS-CoV-2 (Supplementary Fig. 5C). In agreement with a recent study that demonstrated the existence of S mostly in the closed uncleaved pre-fusion confirmation on the authentic SARS-CoV-2 (Turonova et al., 2020), the purified pseudotype SARS-2-S virus spinoculation followed by immunoblotting displayed both glycosylated ∼180 kDa and non-glycosylated ∼120 kDa full-length S0 protein (Fig. 4D). Next, the cells transfected with TMPPSS2 plasmid showed clear processing of pseudotype full-length S0 into S2, and S2’. As expected, pretreatment with Camostat prevented the TMPRSS2 mediated cleavage of pseudotype S0. Since we had observed proteolytic cleavage of ACE2 by TMPRSS2 in the overexpression model (Fig. 3B), we sought to observe whether the TMPRSS2 mediated cleavage of SARS-2-S is taking place in the presence of ACE2. Toward this, we performed immunoprecipitation of TMPRSS2 on pseudotype SARS-2-S spinoculation in the presence or absence of Camostat. Increased pulldown of uncleaved full-length S0 was observed in the Camostat treated cells compared to control, due to the inhibition of TMPRSS2 protease activity and resulting lack of S priming (Fig. 4E). Interestingly, the amount of ACE2 pulled down by TMPRSS2 was not affected by Camostat, suggesting the SARS-2-S cleavage by TMPRSS2 is mediated in the presence of ACE2 in the complex. Noticeably, the cleaved S0 fragments were completely absent in the TMPRSS2 pulldown, suggesting that once processed, the S2 and S2’ fragments are released from the TMPRSS2-ACE2 complex. Similar observations of pseudotype SARS-2-S cleavage by endogenous TMPRSS2 in the presence of endogenous ACE2 were made with LNCaP and Calu-3 cells (Fig. 4F and 4G). These data clearly demonstrate the crucial role played by TMPRSS2 in SARS-CoV-2 viral fusion to the host cell by Spike protein priming (Fig. 4H).

### Camostat alone or in combination with AR directed therapies reduces SARS-2-S pseudovirus entry into prostate and lung cells

After identifying the role of androgen in regulating TMRPSS2 and ACE2 expression, and TMPRSS2 mediated SARS-2-S priming, we asked whether therapeutic intervention targeting AR and TMPRSS2 protease function could affect SARS-CoV-2 infection in prostate and lung cells. Replication-defective virus particles bearing SARS-2-S proteins faithfully reflect critical aspects of SARS-CoV-2 host cell entry (Hoffmann et al., 2020c). Therefore, we employed HIV pseudotypes bearing SARS-2-S and nano-luciferase reporter to test the efficacy of Camostat and AR directed therapies in blocking cell entry (Fig. 5A). We first asked whether androgen deprivation could affect pseudotype virus entry by reducing TMRPSS2 and ACE2 (Fig.2). LNCaP cells grown in the androgen proficient condition were readily susceptible to the entry driven by SARS-2-S (Fig. 5B). However, cells grown under steroid-deprived condition (CSS-charcoal stripped serum) demonstrated a significant reduction in the entry of pseudotype virus. Interestingly, Camostat treatment efficiently blocked the SARS-2-S mediated entry only in the androgen proficient condition, suggesting the presence of TMPRSS2 as a requirement for its potency. Next, we treated LNCaP cells grown in normal growth condition with anti-androgen enzalutamide, AR degrader ARD-69, or Camostat and observed a significant reduction in SARS-2-S driven entry compared to the DMSO control. Notably, the combination of Camostat with enzalutamide or ARD-69 was more efficacious in blocking the entry than single drug (Fig. 5C). Since lungs are the primary target of SARS-CoV-2, we tested whether androgen deprivation, enzalutamide, ARD-69 and Camostat could block SARS-2-S mediated entry in AR positive H460 cells. Similar to LNCaP cells, pseudovirus entry was significantly reduced in the H460 cells grown in the steroid-deprived condition, enzalutamide, and ARD-69 alone or in combination with Camostat compared to the control (Fig. 5D and 5E). As expected, compared to Camostat, the enzalutamide or AR degrader treatment in AR-negative Calu-3 cells did not block the SARS-2-S mediated entry (Fig. 5F). Importantly, unlike SARS-2-S, HIV pseudotypes bearing VSV-G did not display any significant difference in cell entry under these condition in all three tested cell lines (Supplementary Fig. 6). These results indicate that the treatment of AR-positive prostate and lung cells with AR directed therapies in combination with TMPRSS2 inhibitor efficiently block SARS-2-S mediated viral entry.

**Figure 5:**
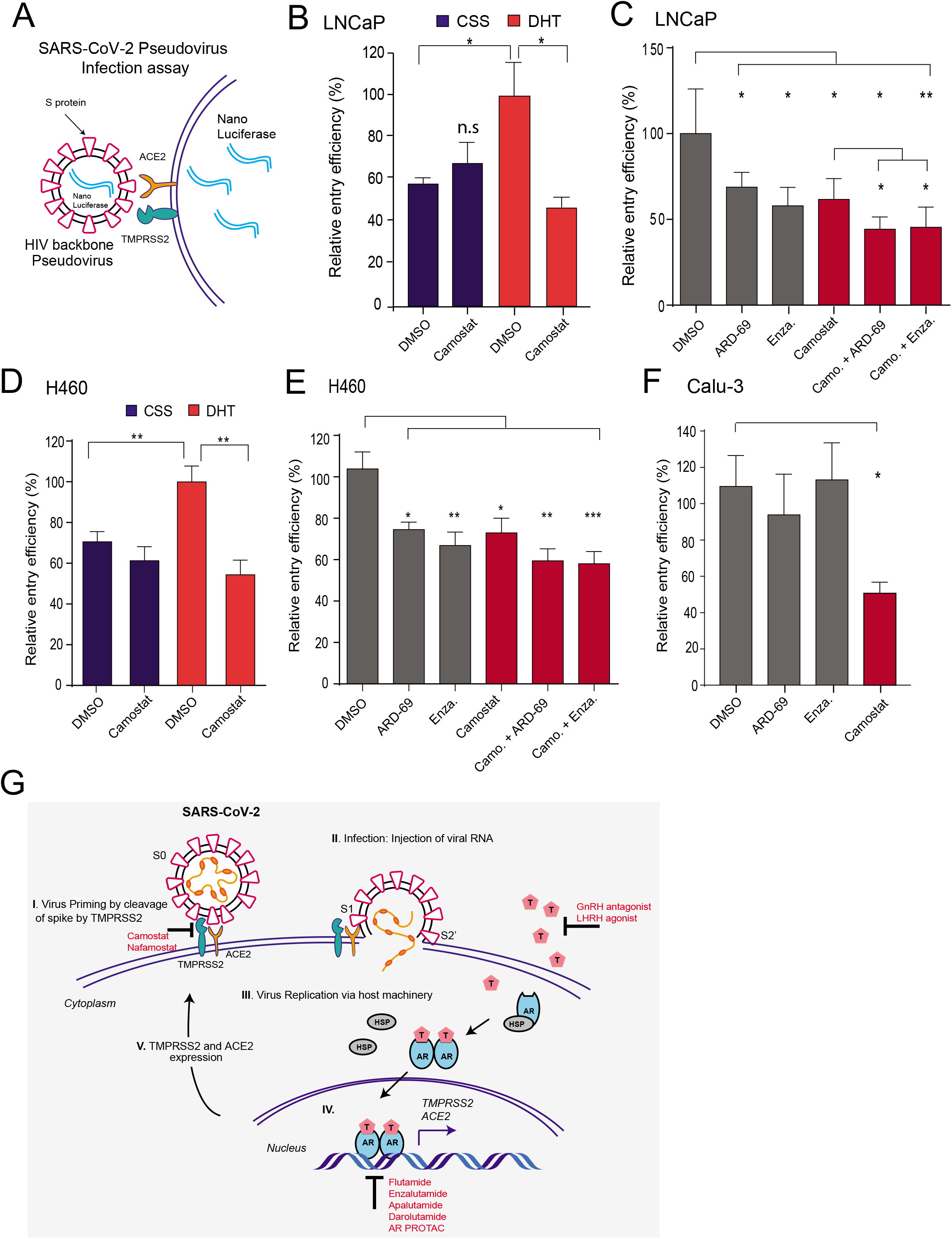
AR targeted therapy in combination with Camostat attenuates the entry of pseudotype SARS-CoV-2 in to the host cells. **(A)** Schematic depicting the SARS-CoV-2 Spike S pseudovirus entry assay. The pseudotype consists of Spike S, and nano-luciferase reporter. **(B)** Androgen deprivation attenuates pseudotype entry. LNCaP cells grown in androgen-deprived serum containing media for three days were pretreated with DMSO, DHT (10 nM) or Camostat (300 μM) for 1h followed by inoculation with SARS-CoV-2 Spike S pseudovirus. 24h post-inoculation, the pseudovirus entry efficiency was measured by means of nano-luciferase signal accompanying entry. The entry efficiency in the DHT treated cells was taken as 100%. Error bar indicates SEM (n =5). **(C)** Anti-androgens or AR degraders with or without Camostat attenuate pseudovirus entry. LNCaP cells grown in complete media was pretreated with enzalutamide (10 μM) or ARD-69 (500 nM) alone, or in combination with Camostat (300 μM) for 1h followed by inoculation with pseudovirus. 24h post-inoculation reporter activity measured as in d. **(D) & (E)**, as in c and d with AR positive H460 lung cells. **(F)** AR negative Calu-3 cells does not respond to anti-androgens. Calu-3 cell are pretreated with enzalutamide, ARD-69 or Camostat followed by pseudovirus inoculation. 24h later reporter signal characterizes the pseudovirus entry efficiency. Error bar indicates SEM (n =5). * P < 0.05, **P< 0.005, ***P< 0.0005 (Student’s t test). **(G)** Schematic depicting the role of TMPRSS2 in SARS-CoV-2 Spike cleavage, and androgen mediated expression of ACE2 and TMPRSS2 that could potentially be targeted by AR directed therapies. Also, see Supplementary Figure 6.

## Discussion

Early intervention of SARS-CoV-2 infection could prevent cytokine storm mediated progression to severe pneumonia and multi-organ failure in COVID-19 patients (Mehta et al., 2020). Although the source of the cytokine storm that causes multi-organ dysfunction is not yet clear, uncontrolled viral replication may contribute to COVID-19 severity and death. The present study provides evidence that the expression of SARS-CoV-2 host cell receptor ACE2 and co-receptor TMPRSS2 is regulated in part by the male sex hormone androgen. Further, our data provide evidence that the cell entry of SARS-CoV-2 Spike pseudovirus can be blocked by androgen deprivation, anti-androgens, or clinically proven inhibitors of TMPRSS2. Although other studies have demonstrated the role of TMRPSS2 in SARS-CoV-2 Spike priming(Hoffmann et al., 2020c), we provide the first direct evidence for the endogenous interaction between TMPRSS2 and ACE2 in human cells, and endogenous TMPRSS2 mediated cleavage of SARS-CoV-2 Spike, which could be blocked by Camostat. This finding is consistent with work demonstrating the role of TMRPSS2 in activating Spike glycoproteins of closely related SARS-CoV and MERS-CoV (Iwata-Yoshikawa et al., 2019; Zhou et al., 2015). Our observation through immunoprecipitation experiments that TMPRSS2-ACE2 exists in complex and potentially required for Spike cleavage has implications for developing novel therapeutics targeting the TMPRSS2-ACE2 interface to block Spike activation. Collectively, our results indicate that androgen regulated TMPRSS2 promotes SARS-CoV-2 entry by two separate mechanisms: ACE2 interaction/cleavage, which might promote viral uptake; and SARS-2-S cleavage, which activates the S protein for membrane fusion (Hoffmann et al., 2020a; Hoffmann et al., 2020c). These results have important implications for our understanding of male bias in COVID-19 severity and mortality, and provide a strong rationale for androgen deprivation or anti-androgen therapy in men with SARS-CoV-2 infection.

The evidence provided by our data and previous studies (Mikkonen et al., 2010) suggests androgen receptor mediated regulation of TMPRSS2 expression in non-prostatic tissues, including lung, may explain the increased susceptibility of men to COVID-19 severity and mortality. Additionally, ACE2 expression has been associated with androgen levels in men, where a lower ACE2 level was associated with older men as a consequence of lower androgen (Chen et al., 2020). Mechanistically, besides androgen, other steroid hormones (estrogen, progesterone, glucocorticoids) may also enhance TMRPSS2 and ACE2 expression in multiple tissues through binding of their respective nuclear receptors to responsive cis elements (ERE, PRE, GRE), which are similar to the ARE present in the gene promoter/enhancer. In addition to the regulation of TMRPSS2 and ACE2 expression, androgen could also increase the SARS-CoV-2 severity in men by modulating the immune response. Androgen is known to increase circulating neutrophils, increase secretion of IL-10, IL-2, IL-8 and TGF-*β* by immune cells, and decrease the antibody response to viral infections(Klein and Flanagan, 2016). It is worth noting that severe cases of COVID-19 exhibit increased neutrophils and IL-8(Barnes et al., 2020; Takahashi et al., 2020).

Except for dexamethasone, which reduces mortality (days alive and free of mechanical ventilation) in ICU patients by reducing inflammation (Tomazini et al., 2020), and Remdesivir, which reduces disease duration by inhibiting viral replication (Goldman et al., 2020), there are currently no drugs for COVID-19. There is an urgent need for prophylactic and therapeutic interventions for patients with existing comorbidities such as cancer as they are at high risk for severe disease due to a weak immune system. Due to an apparent male bias and assumed role of androgen in COVID-19, multiple studies have advocated androgen deprivation therapy (ADT) and anti-androgens – a mainstay of prostate cancer treatment, as potential therapeutic options in COVID-19, which could potentially provide protection against SARS-CoV-2 infection or at least reduce viral amplification (Bhowmick et al., 2020; Mjaess et al., 2020; Montopoli et al., 2020; Patel et al., 2020; Stopsack et al., 2020). Following these assumptions, various clinical trials testing ADT and anti-androgen have begun: Degarelix (GnRH antagonist) NCT04397718, Dutasteride (ADT) NCT04446429, Bicalutamide (first generation anti-androgen) NCT04374279, and Enzalutamide (second-generation anti-androgen) NCT04475601. However, a direct link for androgen regulation of TMPRSS2 and ACE2 in tissues, including the lung, was not established until now. Additionally, Camostat mesilate – the TMPRSS2 inhibitor which is approved for pancreatitis in Japan (Ohshio et al., 1989), is currently being investigated as a treatment for COVID-19 in several active clinical trials in USA (NCT04353284, NCT04470544, NCT04374019), Israel (NCT04355052), and Denmark (NCT04321096). Our data showing complete block of TMRPSS2 mediated Spike activation by Camostat in multiple cell line models provide further credentials for these clinical trials.

Growing evidence suggests cancer patients are susceptible to severe form of COVID-19 mainly due to their immunosuppressive state and co-existing medical conditions (Dai et al., 2020; Lee et al., 2020; Tang and Hu, 2020). In general, patients with metastatic cancers infected with SARS-CoV-2 have had poorer outcomes including death, admission to the intensive care unit, requiring mechanical ventilation, and severe symptoms (Dai et al., 2020). As the development of vaccines and anti-viral drugs against the pandemic causing SARS-CoV-2 are proceeding rapidly, Camostat mesilate along with androgen regulation of TMPRSS2 and ACE2, as a means to inhibit SARS-CoV-2 cell entry (Fig. 5G), and thus, the infection represents a potential strategy in treating COVID-19 in these high-risk population.

### Limitations of the study

Androgen plays a vital role in the human immune response, which we did not address in this paper. Research aimed at characterizing the androgen axis in context with SARS-CoV-2 infection will continue in the future.

## Acknowledgments

We thank Drs. Pöhlmann (UG, Germany), Krogan (UCLA), and Hatziioannou (RU) for SARS-CoV-2 ORFs, nano-luciferase constructs and HEK 293T ACE2 cells. We also thank the Molecular Pathology and Imaging Core (University of Pennsylvania) for histology services, and CHOP Proteomics Core for the mass spectrometry analysis. Research in I.A.A laboratory is supported by NIH (1-R01 CA249210-0), Department of Defense Idea Development Award (W81XWH-17-0404), Conquer Cancer Now Award, and Sarcoma Foundation of America Research Award to I.A.A.

## Author Contributions

I.A.A. conceived the work. I.A.A., Q.D, and R.U., designed the experiments. Q.D., R.U. and R.R performed the experiments and acquired the data. R.N. performed bioinformatics analysis. Q.D, R.U., R.N., and I.A.A., interpreted the data, and prepared the figures. I.A.A. wrote the manuscript with help from all the authors.

## Reference

Global Health 5050. COVID-19 sex-disaggregated data tracker: Sex, gender and COVID-19 (2020).

Afar, D.E., Vivanco, I., Hubert, R.S., Kuo, J., Chen, E., Saffran, D.C., Raitano, A.B., and Jakobovits, A. (2001). Catalytic cleavage of the androgen-regulated TMPRSS2 protease results in its secretion by prostate and prostate cancer epithelia. Cancer Res 61, 1686–1692.

Asangani, I.A., Dommeti, V.L., Wang, X., Malik, R., Cieslik, M., Yang, R., Escara-Wilke, J., Wilder-Romans, K., Dhanireddy, S., Engelke, C., et al. (2014). Therapeutic targeting of BET bromodomain proteins in castration-resistant prostate cancer. Nature 510, 278–282.

Barnes, B.J., Adrover, J.M., Baxter-Stoltzfus, A., Borczuk, A., Cools-Lartigue, J., Crawford, J.M., Dassler-Plenker, J., Guerci, P., Huynh, C., Knight, J.S., et al. (2020). Targeting potential drivers of COVID-19: Neutrophil extracellular traps. J Exp Med 217.

Bhowmick, N.A., Oft, J., Dorff, T., Pal, S., Agarwal, N., Figlin, R.A., Posadas, E.M., Freedland, S.J., and Gong, J. (2020). COVID-19 and androgen-targeted therapy for prostate cancer patients. Endocr Relat Cancer 27, R281–R292.

Cai, Y., Zhang, J., Xiao, T., Peng, H., Sterling, S.M., Walsh, R.M., Jr., Rawson, S., Rits-Volloch, S., and Chen, B. (2020). Distinct conformational states of SARS-CoV-2 spike protein. Science.

Chen, J., Jiang, Q., Xia, X., Liu, K., Yu, Z., Tao, W., Gong, W., and Han, J.J. (2020). Individual variation of the SARS-CoV-2 receptor ACE2 gene expression and regulation. Aging Cell 19.

Consortium, G.T., Laboratory, D.A., Coordinating Center - Analysis Working, G., Statistical Methods groups-Analysis Working, G., Enhancing, G.g., Fund, N.I.H.C., Nih/Nci, Nih/Nhgri, Nih/Nimh, Nih/Nida, et al. (2017). Genetic effects on gene expression across human tissues. Nature 550, 204–213.

Coutard, B., Valle, C., de Lamballerie, X., Canard, B., Seidah, N.G., and Decroly, E. (2020). The spike glycoprotein of the new coronavirus 2019-nCoV contains a furin-like cleavage site absent in CoV of the same clade. Antiviral Res 176, 104742.

Dai, M., Liu, D., Liu, M., Zhou, F., Li, G., Chen, Z., Zhang, Z., You, H., Wu, M., Zheng, Q., et al. (2020). Patients with Cancer Appear More Vulnerable to SARS-CoV-2: A Multicenter Study during the COVID-19 Outbreak. Cancer Discov 10, 783–791.

de Bono, J.S., Guo, C., Gurel, B., De Marzo, A.M., Sfanos, K.S., Mani, R.S., Gil, J., Drake, C.G., and Alimonti, A. (2020). Prostate carcinogenesis: inflammatory storms. Nat Rev Cancer 20, 455–469.

Dubey, R.K., Oparil, S., Imthurn, B., and Jackson, E.K. (2002). Sex hormones and hypertension. Cardiovasc Res 53, 688–708.

Gheblawi, M., Wang, K., Viveiros, A., Nguyen, Q., Zhong, J.C., Turner, A.J., Raizada, M.K., Grant, M.B., and Oudit, G.Y. (2020). Angiotensin-Converting Enzyme 2: SARS-CoV-2 Receptor and Regulator of the Renin-Angiotensin System: Celebrating the 20th Anniversary of the Discovery of ACE2. Circ Res 126, 1456–1474.

Goldman, J.D., Lye, D.C.B., Hui, D.S., Marks, K.M., Bruno, R., Montejano, R., Spinner, C.D., Galli, M., Ahn, M.Y., Nahass, R.G., et al. (2020). Remdesivir for 5 or 10 Days in Patients with Severe Covid-19. N Engl J Med.

Gordon, D.E., Jang, G.M., Bouhaddou, M., Xu, J., Obernier, K., White, K.M., O’Meara, M.J., Rezelj, V.V., Guo, J.Z., Swaney, D.L., et al. (2020). A SARS-CoV-2 protein interaction map reveals targets for drug repurposing. Nature 583, 459–468.

Hamming, I., Timens, W., Bulthuis, M.L., Lely, A.T., Navis, G., and van Goor, H. (2004). Tissue distribution of ACE2 protein, the functional receptor for SARS coronavirus. A first step in understanding SARS pathogenesis. J Pathol 203, 631–637.

Han, X., Wang, C., Qin, C., Xiang, W., Fernandez-Salas, E., Yang, C.Y., Wang, M., Zhao, L., Xu, T., Chinnaswamy, K., et al. (2019). Discovery of ARD-69 as a Highly Potent Proteolysis Targeting Chimera (PROTAC) Degrader of Androgen Receptor (AR) for the Treatment of Prostate Cancer. J Med Chem 62, 941–964.

Henry, G.H., Malewska, A., Joseph, D.B., Malladi, V.S., Lee, J., Torrealba, J., Mauck, R.J., Gahan, J.C., Raj, G.V., Roehrborn, C.G., et al. (2018). A Cellular Anatomy of the Normal Adult Human Prostate and Prostatic Urethra. Cell Rep 25, 3530–3542 e3535.

Heurich, A., Hofmann-Winkler, H., Gierer, S., Liepold, T., Jahn, O., and Pohlmann, S. (2014). TMPRSS2 and ADAM17 cleave ACE2 differentially and only proteolysis by TMPRSS2 augments entry driven by the severe acute respiratory syndrome coronavirus spike protein. J Virol 88, 1293–1307.

Hoffmann, M., Hofmann-Winkler, H., Smith, J.C., Kruger, N., Sorensen, L.K., Sogaard, O.S., Hasselstrom, J.B., Winkler, M., Hempel, T., Raich, L., et al. (2020a). Camostat mesylate inhibits SARS-CoV-2 activation by TMPRSS2-related proteases and its metabolite GBPA exerts antiviral activity. bioRxiv.

Hoffmann, M., Kleine-Weber, H., and Pohlmann, S. (2020b). A Multibasic Cleavage Site in the Spike Protein of SARS-CoV-2 Is Essential for Infection of Human Lung Cells. Mol Cell 78, 779–784 e775.

Hoffmann, M., Kleine-Weber, H., Schroeder, S., Kruger, N., Herrler, T., Erichsen, S., Schiergens, T.S., Herrler, G., Wu, N.H., Nitsche, A., et al. (2020c). SARS-CoV-2 Cell Entry Depends on ACE2 and TMPRSS2 and Is Blocked by a Clinically Proven Protease Inhibitor. Cell 181, 271–280 e278.

Iwata-Yoshikawa, N., Okamura, T., Shimizu, Y., Hasegawa, H., Takeda, M., and Nagata, N. (2019). TMPRSS2 Contributes to Virus Spread and Immunopathology in the Airways of Murine Models after Coronavirus Infection. J Virol 93.

Kawase, M., Shirato, K., van der Hoek, L., Taguchi, F., and Matsuyama, S. (2012). Simultaneous treatment of human bronchial epithelial cells with serine and cysteine protease inhibitors prevents severe acute respiratory syndrome coronavirus entry. J Virol 86, 6537–6545.

Ke, Z., Oton, J., Qu, K., Cortese, M., Zila, V., McKeane, L., Nakane, T., Zivanov, J., Neufeldt, C.J., Cerikan, B., et al. (2020). Structures and distributions of SARS-CoV-2 spike proteins on intact virions. Nature.

Klein, S.L., and Flanagan, K.L. (2016). Sex differences in immune responses. Nat Rev Immunol 16, 626–638.

La Vignera, S., Cannarella, R., Condorelli, R.A., Torre, F., Aversa, A., and Calogero, A.E. (2020). Sex-Specific SARS-CoV-2 Mortality: Among Hormone-Modulated ACE2 Expression, Risk of Venous Thromboembolism and Hypovitaminosis D. Int J Mol Sci 21.

Lee, L.Y.W., Cazier, J.B., Starkey, T., Briggs, S.E.W., Arnold, R., Bisht, V., Booth, S., Campton, N.A., Cheng, V.W.T., Collins, G., et al. (2020). COVID-19 prevalence and mortality in patients with cancer and the effect of primary tumour subtype and patient demographics: a prospective cohort study. Lancet Oncol 21, 1309–1316.

Lucas, J.M., Heinlein, C., Kim, T., Hernandez, S.A., Malik, M.S., True, L.D., Morrissey, C., Corey, E., Montgomery, B., Mostaghel, E., et al. (2014). The androgen-regulated protease TMPRSS2 activates a proteolytic cascade involving components of the tumor microenvironment and promotes prostate cancer metastasis. Cancer Discov 4, 1310–1325.

Mehta, P., McAuley, D.F., Brown, M., Sanchez, E., Tattersall, R.S., Manson, J.J., and Hlh Across Speciality Collaboration, U.K. (2020). COVID-19: consider cytokine storm syndromes and immunosuppression. Lancet 395, 1033–1034.

Mikkonen, L., Pihlajamaa, P., Sahu, B., Zhang, F.P., and Janne, O.A. (2010). Androgen receptor and androgen-dependent gene expression in lung. Mol Cell Endocrinol 317, 14–24.

Millet, J.K., and Whittaker, G.R. (2015). Host cell proteases: Critical determinants of coronavirus tropism and pathogenesis. Virus Res 202, 120–134.

Mjaess, G., Karam, A., Aoun, F., Albisinni, S., and Roumeguere, T. (2020). COVID-19 and the male susceptibility: the role of ACE2, TMPRSS2 and the androgen receptor. Prog Urol.

Montopoli, M., Zumerle, S., Vettor, R., Rugge, M., Zorzi, M., Catapano, C.V., Carbone, G.M., Cavalli, A., Pagano, F., Ragazzi, E., et al. (2020). Androgen-deprivation therapies for prostate cancer and risk of infection by SARS-CoV-2: a population-based study (N = 4532). Ann Oncol 31, 1040–1045.

Montoro, D.T., Haber, A.L., Biton, M., Vinarsky, V., Lin, B., Birket, S.E., Yuan, F., Chen, S., Leung, H.M., Villoria, J., et al. (2018). A revised airway epithelial hierarchy includes CFTR-expressing ionocytes. Nature 560, 319–324.

Moore, J.B., and June, C.H. (2020). Cytokine release syndrome in severe COVID-19. Science 368, 473–474.

Morens, D.M., and Fauci, A.S. (2020). Emerging Pandemic Diseases: How We Got to COVID-19. Cell 182, 1077–1092.

Muus, C., Luecken, M.D., Eraslan, G., Waghray, A., Heimberg, G., Sikkema, L., Kobayashi, Y., Vaishnav, E.D., Subramanian, A., Smilie, C., et al. (2020). Integrated analyses of single-cell atlases reveal age, gender, and smoking status associations with cell type-specific expression of mediators of SARS-CoV-2 viral entry and highlights inflammatory programs in putative target cells. bioRxiv, 2020.2004.2019.049254.

Nersisyan, S., Shkurnikov, M., Turchinovich, A., Knyazev, E., and Tonevitsky, A. (2020). Integrative analysis of miRNA and mRNA sequencing data reveals potential regulatory mechanisms of ACE2 and TMPRSS2. PLoS One 15, e0235987.

Ohshio, G., Saluja, A.K., Leli, U., Sengupta, A., and Steer, M.L. (1989). Esterase inhibitors prevent lysosomal enzyme redistribution in two noninvasive models of experimental pancreatitis. Gastroenterology 96, 853–859.

Patel, V.G., Zhong, X., Liaw, B., Tremblay, D., Tsao, C.K., Galsky, M.D., and Oh, W.K. (2020). Does androgen deprivation therapy protect against severe complications from COVID-19? Ann Oncol.

Pihlajamaa, P., Sahu, B., Lyly, L., Aittomaki, V., Hautaniemi, S., and Janne, O.A. (2014). Tissue-specific pioneer factors associate with androgen receptor cistromes and transcription programs. EMBO J 33, 312–326.

Rasool, R.U., Natesan, R., Deng, Q., Aras, S., Lal, P., Sander Effron, S., Mitchell-Velasquez, E., Posimo, J.M., Carskadon, S., Baca, S.C., et al. (2019). CDK7 Inhibition Suppresses Castration-Resistant Prostate Cancer through MED1 Inactivation. Cancer Discov 9, 1538–1555.

Shang, J., Wan, Y., Luo, C., Ye, G., Geng, Q., Auerbach, A., and Li, F. (2020). Cell entry mechanisms of SARS-CoV-2. Proc Natl Acad Sci U S A 117, 11727–11734.

Shulla, A., Heald-Sargent, T., Subramanya, G., Zhao, J., Perlman, S., and Gallagher, T. (2011). A transmembrane serine protease is linked to the severe acute respiratory syndrome coronavirus receptor and activates virus entry. J Virol 85, 873–882.

Song, H., Seddighzadeh, B., Cooperberg, M.R., and Huang, F.W. (2020). Expression of ACE2, the SARS-CoV-2 Receptor, and TMPRSS2 in Prostate Epithelial Cells. Eur Urol 78, 296–298.

Stopsack, K.H., Mucci, L.A., Antonarakis, E.S., Nelson, P.S., and Kantoff, P.W. (2020). TMPRSS2 and COVID-19: Serendipity or Opportunity for Intervention? Cancer Discov 10, 779–782.

Takahashi, T., Ellingson, M.K., Wong, P., Israelow, B., Lucas, C., Klein, J., Silva, J., Mao, T., Oh, J.E., Tokuyama, M., et al. (2020). Sex differences in immune responses that underlie COVID-19 disease outcomes. Nature.

Tang, L.V., and Hu, Y. (2020). Poor clinical outcomes for patients with cancer during the COVID-19 pandemic. Lancet Oncol 21, 862–864.

Tomazini, B.M., Maia, I.S., Cavalcanti, A.B., Berwanger, O., Rosa, R.G., Veiga, V.C., Avezum, A., Lopes, R.D., Bueno, F.R., Silva, M., et al. (2020). Effect of Dexamethasone on Days Alive and Ventilator-Free in Patients With Moderate or Severe Acute Respiratory Distress Syndrome and COVID-19: The CoDEX Randomized Clinical Trial. JAMA.

Tomlins, S.A., Rhodes, D.R., Perner, S., Dhanasekaran, S.M., Mehra, R., Sun, X.W., Varambally, S., Cao, X., Tchinda, J., Kuefer, R., et al. (2005). Recurrent fusion of TMPRSS2 and ETS transcription factor genes in prostate cancer. Science 310, 644–648.

Turonova, B., Sikora, M., Schurmann, C., Hagen, W.J.H., Welsch, S., Blanc, F.E.C., von Bulow, S., Gecht, M., Bagola, K., Horner, C., et al. (2020). In situ structural analysis of SARS-CoV-2 spike reveals flexibility mediated by three hinges. Science.

Vabret, N., Britton, G.J., Gruber, C., Hegde, S., Kim, J., Kuksin, M., Levantovsky, R., Malle, L., Moreira, A., Park, M.D., et al. (2020). Immunology of COVID-19: Current State of the Science. Immunity 52, 910–941.

Wadman, M. (2020). Sex hormones signal why virus hits men harder. Science 368, 1038–1039.

Walls, A.C., Park, Y.J., Tortorici, M.A., Wall, A., McGuire, A.T., and Veesler, D. (2020). Structure, Function, and Antigenicity of the SARS-CoV-2 Spike Glycoprotein. Cell 181, 281–292 e286.

Walls, A.C., Tortorici, M.A., Snijder, J., Xiong, X., Bosch, B.J., Rey, F.A., and Veesler, D. (2017). Tectonic conformational changes of a coronavirus spike glycoprotein promote membrane fusion. Proc Natl Acad Sci U S A 114, 11157–11162.

Wang, C., Horby, P.W., Hayden, F.G., and Gao, G.F. (2020). A novel coronavirus outbreak of global health concern. Lancet 395, 470–473.

Williamson, E.J., Walker, A.J., Bhaskaran, K., Bacon, S., Bates, C., Morton, C.E., Curtis, H.J., Mehrkar, A., Evans, D., Inglesby, P., et al. (2020). Factors associated with COVID-19-related death using OpenSAFELY. Nature 584, 430–436.

Zhang, X., Tan, Y., Ling, Y., Lu, G., Liu, F., Yi, Z., Jia, X., Wu, M., Shi, B., Xu, S., et al. (2020). Viral and host factors related to the clinical outcome of COVID-19. Nature 583, 437–440.

Zhou, Y., Vedantham, P., Lu, K., Agudelo, J., Carrion, R., Jr., Nunneley, J.W., Barnard, D., Pohlmann, S., McKerrow, J.H., Renslo, A.R., et al. (2015). Protease inhibitors targeting coronavirus and filovirus entry. Antiviral Res 116, 76–84.

Ziegler, C.G.K., Allon, S.J., Nyquist, S.K., Mbano, I.M., Miao, V.N., Tzouanas, C.N., Cao, Y., Yousif, A.S., Bals, J., Hauser, B.M., et al. (2020). SARS-CoV-2 Receptor ACE2 Is an Interferon-Stimulated Gene in Human Airway Epithelial Cells and Is Detected in Specific Cell Subsets across Tissues. Cell 181, 1016–1035 e1019.

