## supplementary figures and methods for "Targeting androgen regulation of TMPRSS2 and ACE2 as a therapeutic strategy to combat COVID-19"

### Supplementary Figure 1

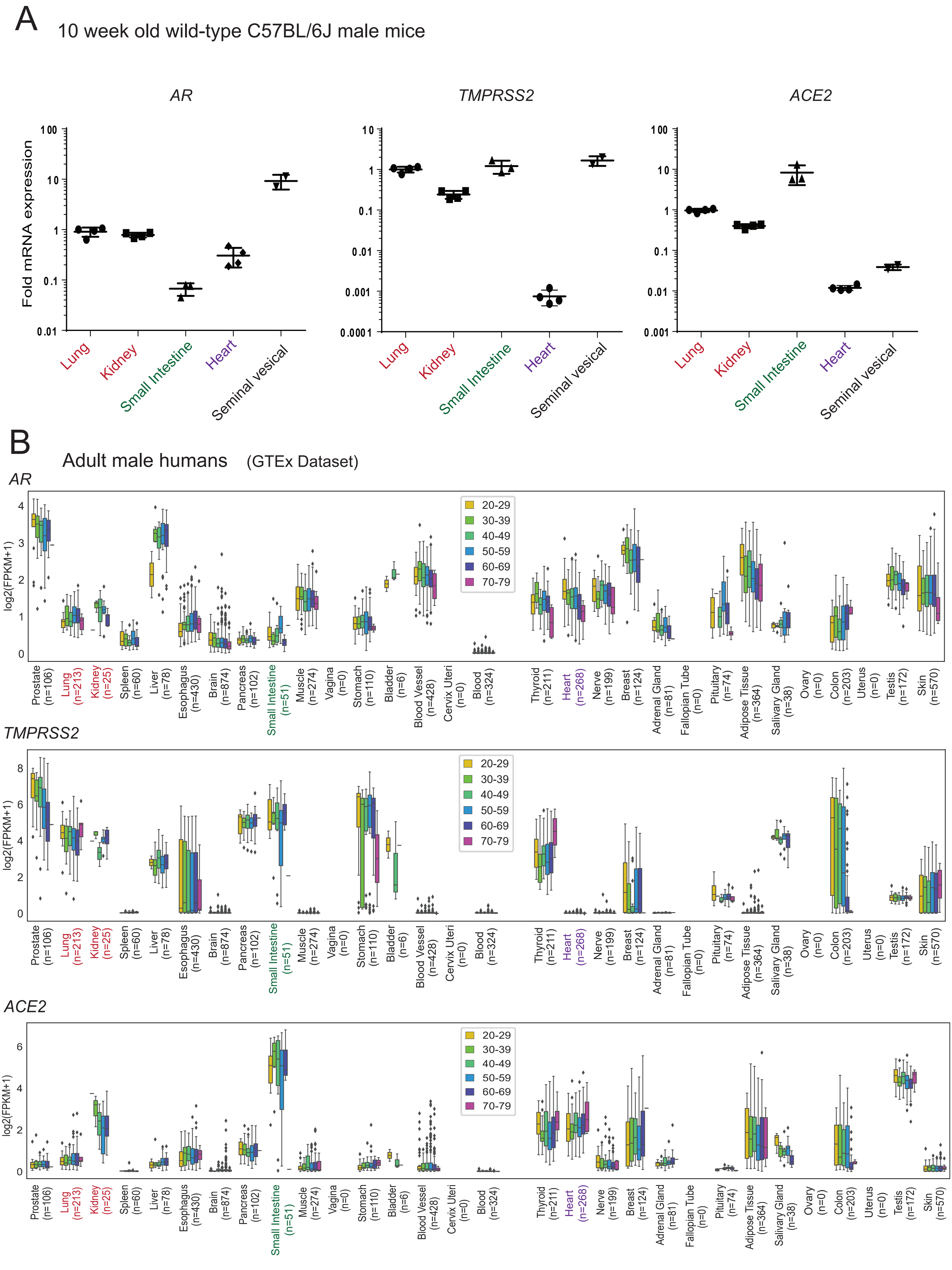

**Supplementary Figure 1. Expression of AR, ACE2 and TMPRSS2 in mice and human male.** **(A)** Expression of AR, ACE2 and TMPRSS2 in various organs in adult male mice. qRT-PCR analysis for AR in the indicated organs from mock castrated male mice. **(B)** GTEx data showing the AR, TMPRSS2 and ACE2 mRNA expression in various organs in human male. Related to Figure. 1.

### Supplementary Figure 2

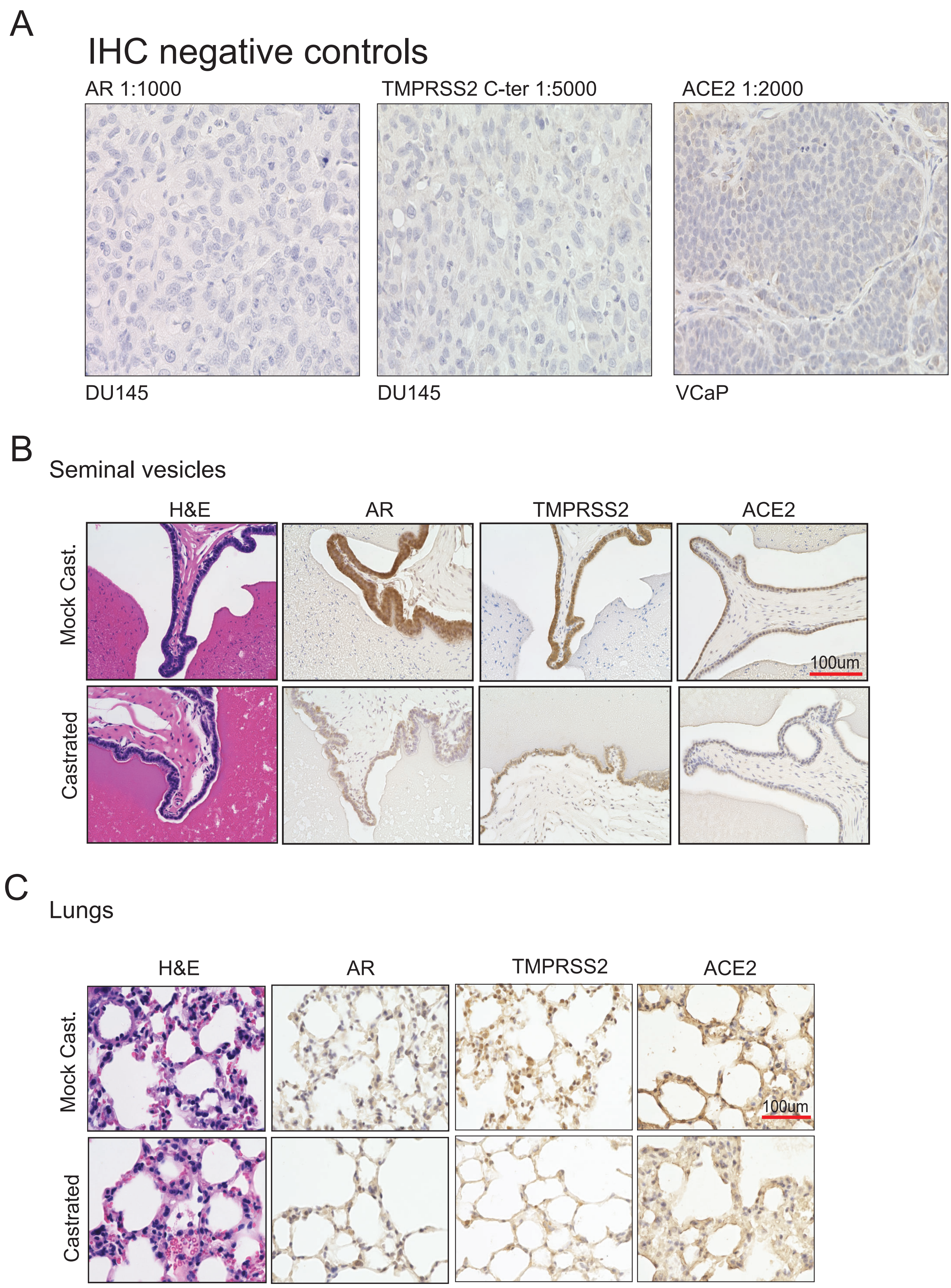

**Supplementary Figure 2. Immunohistochemistry staining for AR, TMPRSS2 and ACE2 in mice and human cells. (A)** Negative control for immunohistochemistry staining data presented in Figure 1D and 1E. Immunohistochemistry with AR, TMRPSS2 and ACE2 antibody at the indicated dilution was performed on AR and TMPRSS2 negative DU145 xenograft tissue section, and ACE2 negative VCaP xenograft tissue section. **(B)** Immunohistochemistry analysis of the indicated target protein in the seminal vesicles from mock and castrated males. Note the reduced AR, TMPRSS2, and ACE2 staining in the castrated group compared to mock. **(C)** Higher magnification images for the staining of lung tissues shown in Fig. 1E. Note the reduced TMPRSS2 and ACE2 staining intensity in the castrated group. Related to Figure. 1.

### Supplementary Figure 3

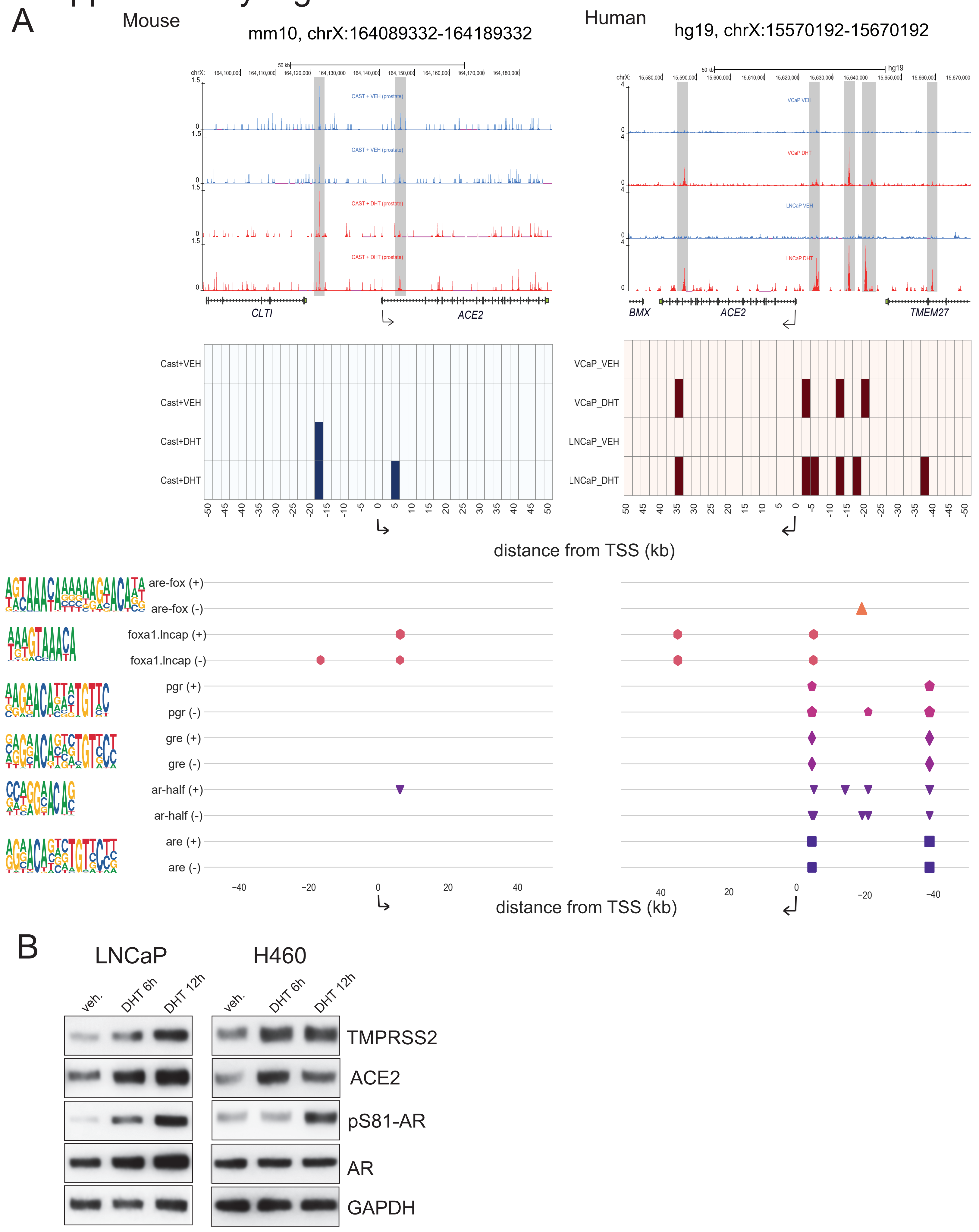

**Supplementary Figure 3.** ***TMPRSS2*** **and** ***ACE2*** **loci displays enhanced AR-binding upon testosterone stimulation. (A)** Shown are AR ChIP-seq tracks in vehicle and testosterone treated conditions, their enrichment profile, and the locations of six canonical AR motifs, in the region 50 kb up- and down-stream of *Ace2* (left panel) and *ACE2* (right panel). The shown tracks are from the publicly available datasets GSE47192 (castrated mouse prostate) and GSE125245 (VCaP and LNCaP lines previously published from our laboratory). All dataset were analyzed as described in the methods section. **(B)** Increase in TMPRSS2 and ACE2 protein upon DHT stimulation. LNCaP and H460 cells were grown in CSS media for 48h followed by stimulation with 10 nM for 6h and 12h. Total lysates prepared were used for immunoblotting with indicated antibodies. GAPDH was used loading control. Related to Figure. 2.

### Supplementary Figure 4

A

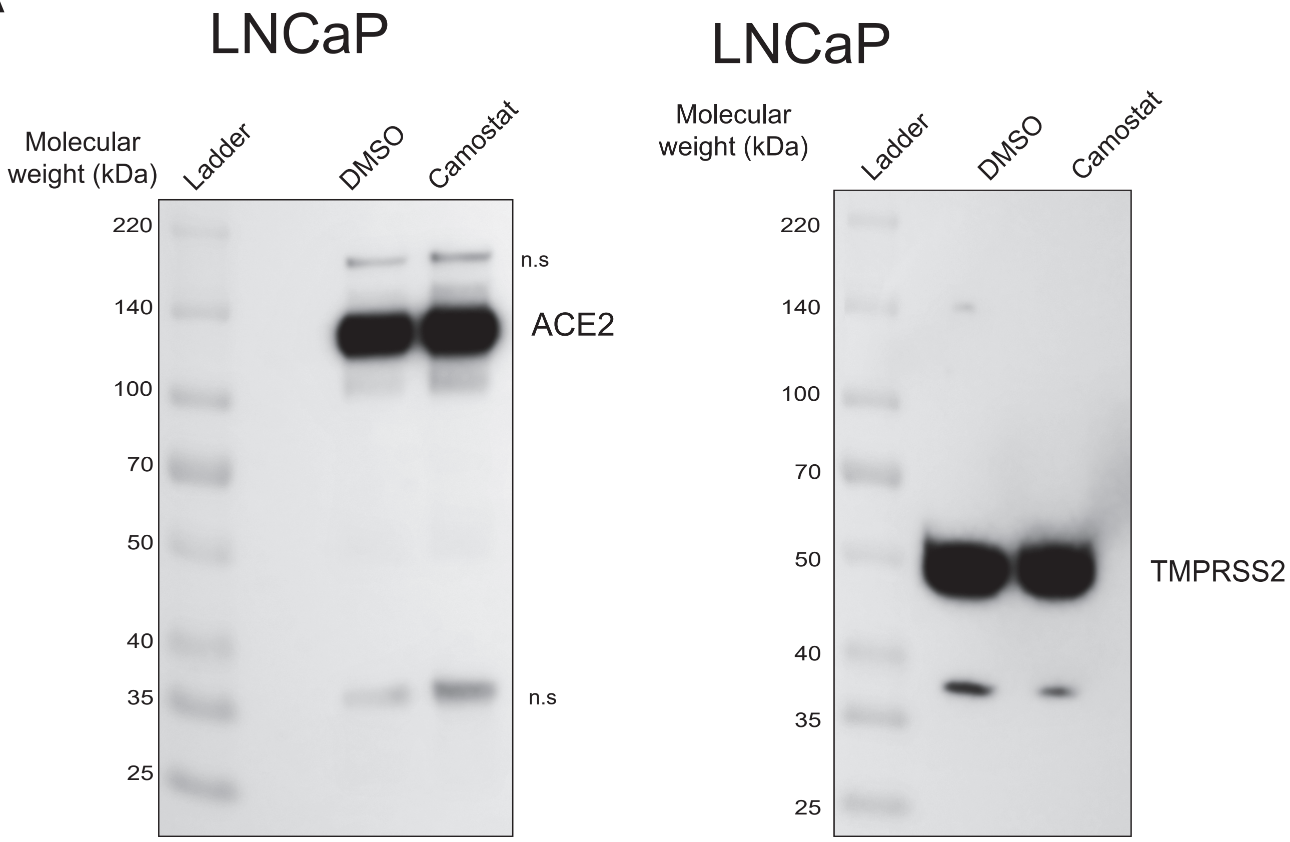


B

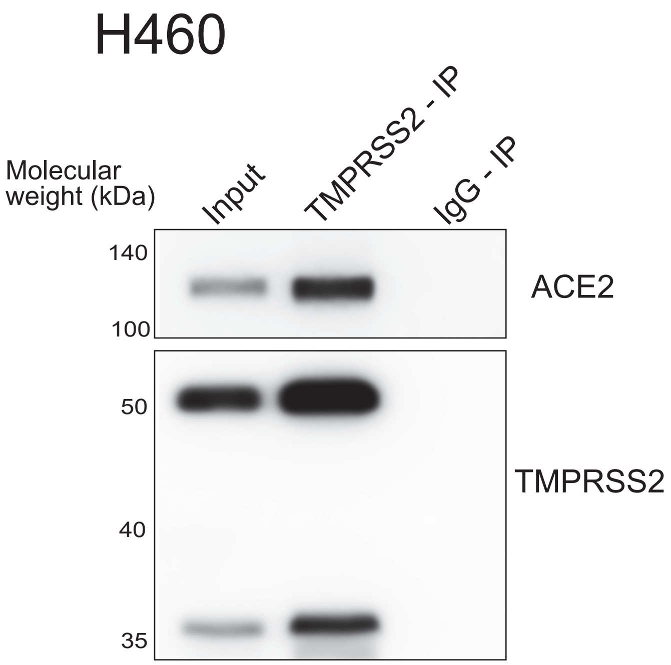

**Supplementary Figure 4. Effect of Camostat on ACE2, and interaction between ACE2-TMPRSS2 in H460 cells. (A) Camostat does not affect TMRPSS2 or ACE2 levels.** Immunoblot showing TMPRSS2and ACE2 levels in LNCaP cells treated with DMSO or Camostat (250 uM) for 24h. Protein molecular weight ladder used to determine the exact size of the target protein. **(B)** TMPRSS2 and ACE2 physically interacts in H460 cells. Immunoprecipitation using C-terminal TMPRSS2 antibody with H460 protein extracts, followed by immunoblotting with ACE2 and TMRPSS2 antibody. IgG pulldown served as a negative control; input was 10%. Related to Figure. 3.

### Supplementary Figure 5

A

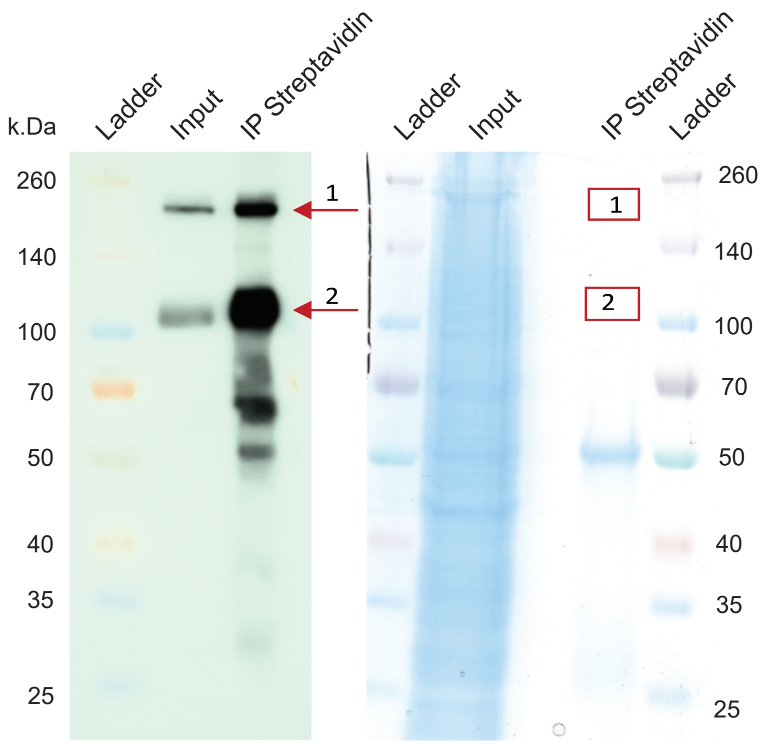


C

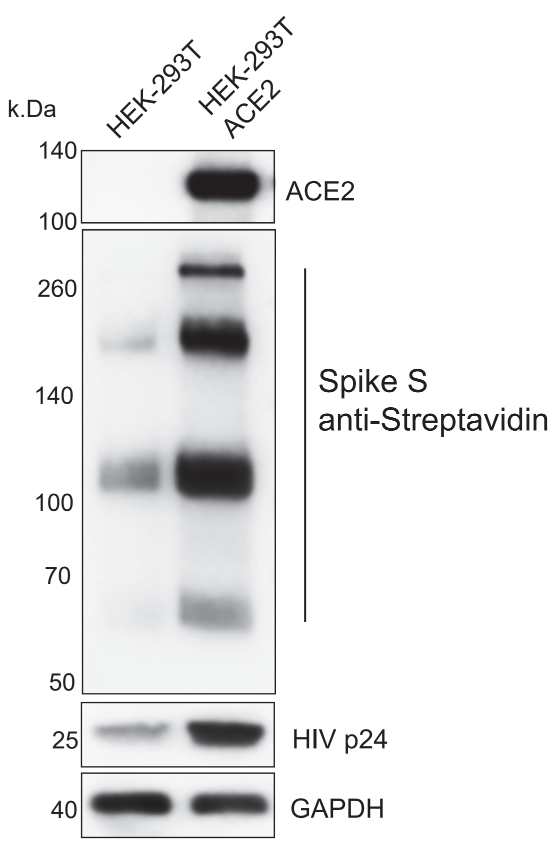


B

#### Band 1

**P0DTC2 (100%), 141,180.7 Da**
**Spike glycoprotein OS=Severe acute respiratory syndrome coronavirus 2 OX=2697049 GN=S PE=1 SV=1**
**52 exclusive unique peptides, 78 exclusive unique spectra, 210 total spectra, 654/1273 amino acids (51% coverage)**

MFVFLVLLPL VSSQCVNLTT RTQLPPAYTN SFTRGVYYPD KVFRSSVLHS TQDLFLPFFS NVTWFHAIHV SGTNGTKRFD
NPVLPFNDGV YFASTEKSNI IRGWIFGTTL DSKTQSLLIV NNATNVVIKV CEFQFCNDPF LGVYYHKNNK SWMESEFRVY
SSANNCTFEY VSQPFLMDLE GKQGNFKNLR EFVFKNIDGY FKIYSKHTPI NLVRDLPQGF SALEPLVDLP IGINITRFQT
LLALHRSYLT PGDSSSGWTA GAAAYYVGYL QPRTFLLKYN ENGTITDAVD CALDPLSETK CTLKSFTVEK GIYQTSNFRV
QPTESIVRFP NITNLCPFGE VFNATRFASV YAWNRKRISN CVADYSVLYN SASFSTFKCY GVSPTKLNDL CFTNVYADSF
VIRGDEVRQI APGQTGKIAD YNYKLPDDFT GCVIAWNSNN LDSKVGGNYN YLYRLFRKSN LKPFERDIST EIYQAGSTPC
NGVEGFNCYF PLQSYGFQPT NGVGYQPYRV VVLSFELLHA PATVCGPKKS TNLVKNKCVN FNFNGLTGTG VLTESNKKFL
PFQQFGRDIA DTTDAVRDPQ TLEILDITPC SFGGVSVITP GTNTSNQVAV LYQDVNCTEV PVAIHADQLT PTWRVYSTGS
NVFQTRAGCL IGAEHVNNSY ECDIPIGAGI CASYQTQTNS PRRARSVASQ SIIAYTMSLG AENSVAYSNN SIAIPTNFTI
SVTTEILPVS MTKTSVDCTM YICGDSTECS NLLLQYGSFC TQLNRALTGI AVEQDKNTQE VFAQVKQIYK TPPIKDFGGF
NFSQILPDPS KPSKRSFIED LLFNKVTLAD AGFIKQYGDC LGDIAARDLI CAQKFNGLTV LPPLLTDEMI AQYTSALLAG
TITSGWTFGA GAALQIPFAM QMAYRFNGIG VTQNVLYENQ KLIANQFNSA IGKIQDSLSS TASALGKLQD VVNQNAQALN
TLVKQLSSNF GAISSVLNDI LSRLDKVEAE VQIDRLITGR LQSLQTYVTQ QLIRAAEIRA SANLAATKMS ECVLGQSKRV
DFCGKGYHLM SFPQSAPHGV VFLHVTYVPA QEKNFTTAPA ICHDGKAHFP REGVFVSNGT HWFVTQRNFY EPQIITTDNT
FVSGNCDVVI GIVNNTVYDP LQPELDSFKE ELDKYFKNHT SPDVDLGDIS GINASVVNIQ KEIDRLNEVA KNLNESLIDL
QELGKYEQYI KWPWYIWLGF IAGLIAIVMV TIMLCCMTSC CSCLKGCCSC GSCCKFDEDD SEPVLKGVKL HYT

#### Band 2

**P0DTC2 (100%), 141,180.7 Da**
**Spike glycoprotein OS=Severe acute respiratory syndrome coronavirus 2 OX=2697049 GN=S PE=1 SV=1**
**57 exclusive unique peptides, 82 exclusive unique spectra, 224 total spectra, 682/1273 amino acids (54% coverage)**

MFVFLVLLPL VSSQCVNLTT RTQLPPAYTN SFTRGVYYPD KVFRSSVLHS TQDLFLPFFS NVTWFHAIHV SGTNGTKRFD
NPVLPFNDGV YFASTEKSNI IRGWIFGTTL DSKTQSLLIV NNATNVVIKV CEFQFCNDPF LGVYYHKNNK SWMESEFRVY
SSANNCTFEY VSQPFLMDLE GKQGNFKNLR EFVFKNIDGY FKIYSKHTPI NLVRDLPQGF SALEPLVDLP IGINITRFQT
LLALHRSYLT PGDSSSGWTA GAAAYYVGYL QPRTFLLKYN ENGTITDAVD CALDPLSETK CTLKSFTVEK GIYQTSNFRV
QPTESIVRFP NITNLCPFGE VFNATRFASV YAWNRKRISN CVADYSVLYN SASFSTFKCY GVSPTKLNDL CFTNVYADSF
VIRGDEVRQI APGQTGKIAD YNYKLPDDFT GCVIAWNSNN LDSKVGGNYN YLYRLFRKSN LKPFERDIST EIYQAGSTPC
NGVEGFNCYF PLQSYGFQPT NGVGYQPYRV VVLSFELLHA PATVCGPKKS TNLVKNKCVN FNFNGLTGTG VLTESNKKFL
PFQQFGRDIA DTTDAVRDPQ TLEILDITPC SFGGVSVITP GTNTSNQVAV LYQDVNCTEV PVAIHADQLT PTWRVYSTGS
NVFQTRAGCL IGAEHVNNSY ECDIPIGAGI CASYQTQTNS PRRARSVASQ SIIAYTMSLG AENSVAYSNN SIAIPTNFTI
SVTTEILPVS MTKTSVDCTM YICGDSTECS NLLLQYGSFC TQLNRALTGI AVEQDKNTQE VFAQVKQIYK TPPIKDFGGF
NFSQILPDPS KPSKRSFIED LLFNKVTLAD AGFIKQYGDC LGDIAARDLI CAQKFNGLTV LPPLLTDEMI AQYTSALLAG
TITSGWTFGA GAALQIPFAM QMAYRFNGIG VTQNVLYENQ KLIANQFNSA IGKIQDSLSS TASALGKLQD VVNQNAQALN
TLVKQLSSNF GAISSVLNDI LSRLDKVEAE VQIDRLITGR LQSLQTYVTQ QLIRAAEIRA SANLAATKMS ECVLGQSKRV
DFCGKGYHLM SFPQSAPHGV VFLHVTYVPA QEKNFTTAPA ICHDGKAHFP REGVFVSNGT HWFVTQRNFY EPQIITTDNT
FVSGNCDVVI GIVNNTVYDP LQPELDSFKE ELDKYFKNHT SPDVDLGDIS GINASVVNIQ KEIDRLNEVA KNLNESLIDL
QELGKYEQYI KWPWYIWLGF IAGLIAIVMV TIMLCCMTSC CSCLKGCCSC GSCCKFDEDD SEPVLKGVKL HYT

**Supplementary Figure 5. Mass spectrometry analysis of pseudotype SARS-2-S proteins. (A)**, *Left,* Proteins extracted from purified pseudotype SARS-2-S virus were subjected to immunoprecipitation with streptavidin antibody followed by immunoblotting. Arrow indicates two prominent bands analyzed for sequencing. *Right,* SDS-PAGE and Coomassie stain gel with the streptavidin immunoprecipitated eluates. Box indicates the excised regions of gel used for protein extraction for mass spectrometry. **(B)** SARS-2-S protein sequence with peptide coverage highlighted in yellow for band 1 and band 2. Total spectra and unique peptides for both the bands are indicated on the top. **(C)** Pseudotype SARS-CoV-2 Spike requires ACE2 for cell entry. HEK 293T and HEK 293T ACE2 overexpressing cells were infected with pseudotype HIV bearing SARS-CoV-2 Spike via spinoculation for 1h. Next, 4h post-spinoculation recovery total lysates were prepared and used for immunoblotting with the indicated antibody. Note the increased signal for different forms of Spike proteins and HIV p24 in the ACE2 expressing cells. GAPDH was used as a loading control. Related to Figure. 4.

### Supplementary Figure 6

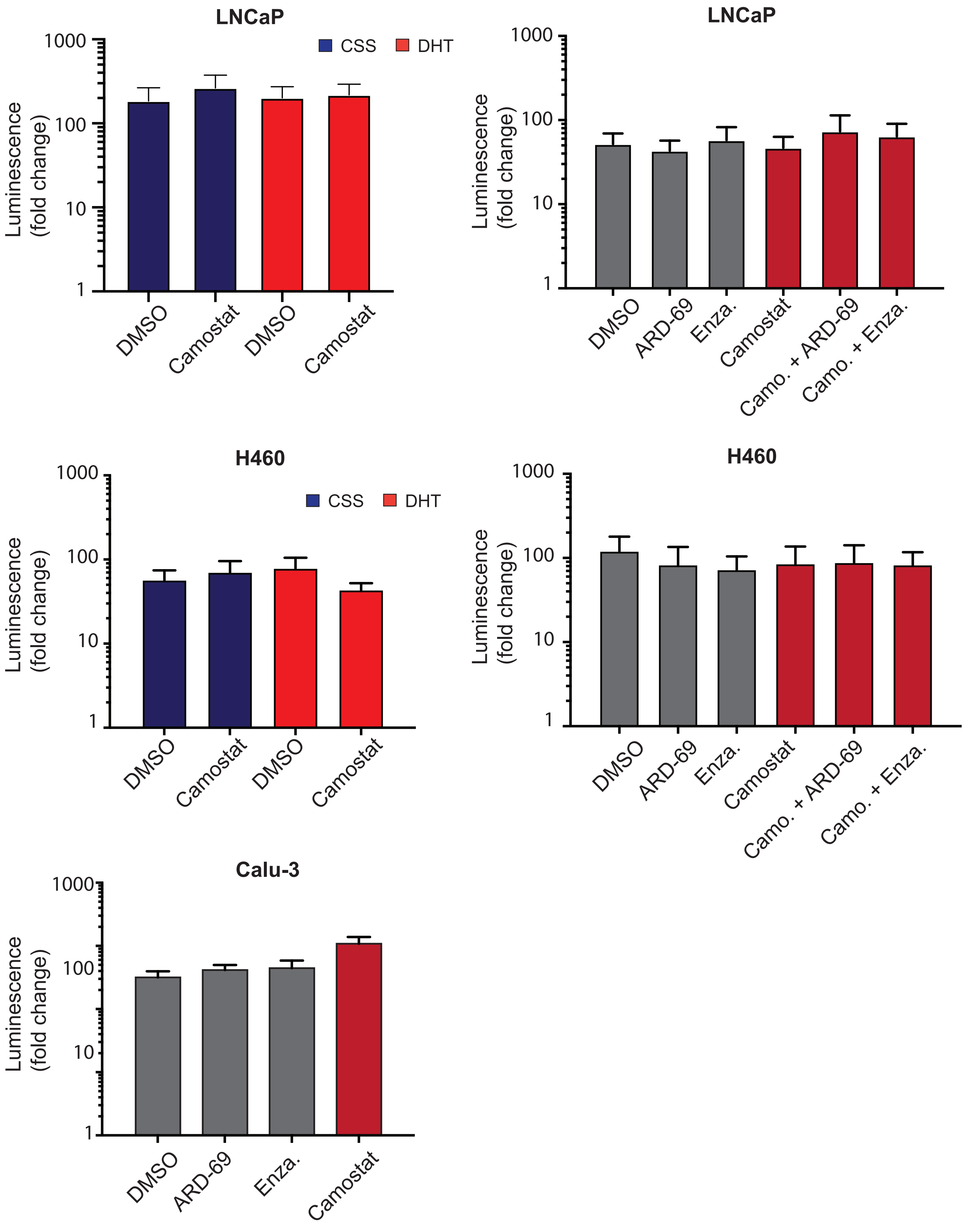

**Supplementary Figure 6. VSV-G pseudotype entry is not affected by ADT, anti-androgens or Camostat treatment.** As in figure 5 with VSV-G pseudotype entry in LNCaP, H460 and Calu-3 cells under different conditions as in Figure 5. Briefly, LNCaP and H460 cells grown in androgen-deprived serum containing media for three days were pretreated with DMSO, DHT (10 nM) or Camostat (300 μM) for 1h followed by inoculation with VSV-G pseudovirus. 24h post-inoculation, the pseudovirus entry efficiency was measured by means of nano-luciferase signal accompanying entry. LNCaP and H460 cells grown in complete media was pretreated with enzalutamide (10 μM) or ARD-69 (500 nM) alone, or in combination with Camostat (300 μM) for 1h followed by inoculation with VSV-G pseudovirus. 24h post-inoculation reporter activity was measured. Calu-3 cell were pretreated with enzalutamide**,** ARD-69 or Camostat followed by VSV-G pseudovirus inoculation. 24h later reporter signal characterizes the pseudovirus entry efficiency. Error bar indicates SEM (n =5). Related to Figure. 5.

#### Transparent Methods

##### Mouse castration experiment

*In vivo* mouse studies were performed following protocols approved by the Institutional Animal Care and Use Committee at the University of Pennsylvania and in compliance with all regulatory standards. C57BL/6J mice were procured from Jackson Laboratory (000664) and breed in house. Eight-to-nine-week-old C57BL/6J male mice were randomized into two groups; one group underwent surgical castration, while the other group had a mock surgery. 8 days post-castration, mice were euthanized, and major organs were harvested for RNA, protein, and immunohistochemistry analysis. The female mice were used as control.

##### Immunohistochemistry

Tissues were fixed in 4% of formaldehyde for 48h, and paraffin embedded through Molecular Pathology and Imaging Core at UPENN. Paraffin embedded sections from the tissue of mock castrated male, castrated male and female mice were deparaffinized with 3 changes of xylene for 5 min each. Slides were then rehydrated in 100% alcohol for 10min, 95% alcohol 10min for 2 changes, 70% alcohol and distilled water for 10min each. Slides were subjected to citrate based (pH 6.0) antigen retrieval (Vector, H-3300) at 95°C for 30mins and followed by blocking endogenous peroxidase activity with 3% $H_2O_2$ for 5min. After three times of 5 min TBST washing, slides were with blocking buffer (1.25% of goat serum) at room temperature for one hour. Slides were applied with diluted primary antibody and incubated in a humidified chamber at 4°C overnight. After three times of 10min TBST washing, slides were incubated with secondary antibody (Vector, PK-4001 and R&D, CTS008) for 30min at room temperature. After three times of 10min TBST washing, the color of the antibody staining was revealed by peroxidase-based detection (R&D, CTS008) and the sections were counterstained with hematoxylin (Millipore Sigma, MHS1). Representative photographs were taken on a Keyence BZ-X Series All-in-one Fluorescence Microscope with a 20x and 40 x objectives. Images were assessed and quantified with ImageJ.

##### ChIP-seq analysis

Publically available castrated mouse prostate AR ChIP-seq (Chromatin Immunoprecipitation followed by sequencing) data submitted to Gene Expression Omnibus (GSE47192) was downloaded as raw fastq files, quality checked using FASTQC (www.bioinformatics.babraham.ac.uk/projects/fastqc) and aligned to the GRCm38 (release 19) genome using the STAR v2.7.3a aligner with default settings. PCR duplicate reads in the aligned bam files were removed using samtools, bam files were converted to CPM normalized bigwig tracks using deeptools, and viewed using the Integrated Genomics Viewer or the UCSC genome browser. Single-end reads were extended up to the fragment length (200 bp) along the read direction. ChIP-sequencing data for human prostate lines VCaP and LNCaP from our lab and previously submitted to GEO (GSE125245) were analyzed as described and aligned to GRCh37 (release 10) reference genome.

**ChIP-Enrichment analysis**

AR enrichment peaks in both human and mouse ChIP-seq data were computed using MACS2(Zhang et al., 2008), with default settings.  Regions found to ubiquitously enriched across a number of next-generation sequencing experiments, also known as the blacklisted peaks (https://sites.google.com/site/anshulkundaje/projects/blacklists), were excluded in all subsequent analysis. Heatmaps of various ChIP-enriched regions (2500 kbp spatial resolution, p-value<0.05) were assembled using in-house python scripts.

**Motif enrichment analysis**

The presence and enrichment of various AR-binding motifs within AR-enriched regions 50 kb up- and down-stream of *ACE2*/*Ace2* were determined using HOMER(Heinz et al., 2010). Briefly, position weight matrices for well-established AR motifs, namely **are, are-fox, ar-half, foxa1.lncap, gre,** and **pgr** (all sourced from the HOMER package), were supplied to findGenomeMotifs.pl using the -find option. The resulting enrichment data was plotted as a scatter plot as shown in Supplementary Fig. 2A using in-house python scripts.

**Gene expression analysis**

The transcriptomic dataset from Genotype-Tissue Expression (GTEx V6p, n = 11401) and associated metadata (Consortium et al., 2017) were downloaded from GTEx website and analyzed using in-house scripts. The metadata was incorporated into the expression matrix to construct sex, age, and organ-specific cohorts for *AR, ACE2* and *TMPRSS2* as shown in Supplementary Figure 1B.

**RNA extraction and quantitative RT-PCR**

Total RNA was isolated from cells and tissue using miRNeasy Mini Kit (Qiagen,74106), and cDNA was synthesized from 1μg total RNA using SuperScriptIV (Life Technologies,18090200). qRT-PCR was performed in triplicates using Fast SYBR Master Mix (Life Technologies, 4385617), and analyzed on QuantStudio3 (Applied Biosystems, USA). The target mRNA expression was quantified using ΔΔCt method, as normalized to GAPDH transcript levels. Primers were designed using Primer3 Input (version 0.4.0) (http://bioinfo.ut.ee/primer3-0.4.0/primer3) and synthesized by Integrated DNA Technologies. See supplementary table 1 for the primer sequence.

**Cell culture**

Prostate cancer cell line LNCaP (obtained from ATCC) and lung cancer cell lines H838 and H460 (obtained from ATCC) were maintained in RPMI-1640 media (Gibco, 11875093). Lung cancer cell line Calu-3 (obtained from ATCC) was maintained in DMEM/F-12 (Gibco, 11320033). HEK-293T cells (obtained from ATCC) and HEK-293T-ACE2cl.22 (Gift from Paul D. Bieniasz, Rockefeller University) were maintained in DMEM media (Gibco, 21013024). The medium was supplemented with 10% of Fetal Bovine Serum (FBS) (HYC, SH30910.03) and 1% of Penicillin Streptomycin Solution (Invitrogen, 15140122), hereto referred to as "complete media". All cell lines were tested negative for mycoplasma contamination, and were maintained in a humidified incubator at 37°C and 5% $CO_2$.

**Drugs/Chemicals**

The enzalutamide (Selleckchem, S1250), Camostat Mesylate (Selleck Chemicals, 59721-29-8) and PROTAC-ARD-69 (gift from Dr. Shaomeng Wang, University of Michigan, Ann Arbor, MI, Ref: http://dx.doi.org/10.1021/acs.jmedchem.8b01631) were dissolved and aliquoted in DMSO (Sigma-Aldrich, D2650). 5α-Dihydrotestosterone (Cerilliant, D073) was dissolved and aliquoted in methanol.

**Androgen deprivation and Drug treatments**

For starvation or DHT stimulation for AR signaling, LNCaP, and H460 cells were plated in complete media at required confluency. After 24h cells were washed with DPBS and cultured in RPMI 1640 without Phenol Red (Invitrogen-11835030) with 10% charcoal-dextran stripped FBS (Gemini Bio-Products- 100119) for 2, 4 and 6 days. Regular RPMI 1640 medium with 10% FBS was included as a control. For drug treatments, the cells after 24h of seeding in complete media were exposed to either 25 μM Enzalutamide or 250 nM PROTAC ARD-69 for 3 and 6 days. Cells treated with DMSO were used as a control. After the treatment time points, the total RNA and protein lysates were prepared for qRT-PCR and immunoblotting, respectively.

**Immunoblotting and Co-immunoprecipitation**

Cell lysates were prepared with RIPA buffer (Boston Bioproducts, BP-115DG)-10 mM Tris-HCl pH 7.5, 1 mM EDTA, 400 mM NaCl, 0.5% NP-40, and 1 mM DTT, supplemented with protease inhibitor cocktail (Pierce, A32965) and phosphatase inhibitor (Thermo, 1861280). For immunoblotting, the total protein lysates were boiled at 100°C in Laemmli Sample Buffer (Bio-rad, 1610737) and then separated by SDS-PAGE and transferred onto Polyvinylidene Difluoride membrane (GE Healthcare, IPVH00010). The membrane was incubated for one hour in blocking buffer [Tris-buffered saline, 0.1% Tween (TBS-T), 5% nonfat dry milk] followed by overnight incubation on a rocker at 4°C with the primary antibody. Following a wash with TBS-T, the membrane was incubated with horseradish peroxidase-conjugated secondary antibody for 2h on a rocker at room temperature. The membrane was washed again, and signals were visualized using enhanced chemiluminescence system as per manufacturer's protocol (GE Healthcare) or Kwik Quant Imager (Kindle Biosciences). All antibodies were employed at the dilutions suggested by the manufacturers.

For immunoprecipitation experiments, protein extracts were obtained from cells using IP buffer (20 mM Tris pH7.5, 150 mM NaCl, 1% Triton-X 100, 5mM EDTA, Protease/Phosphatase Inhibitor) by sonication. The lysates (0.35-1.0 mg) were pre-cleaned by incubation with protein G Dyna beads (Thermo Fisher Scientific, 10004D) for 1h on a rotator at 4°C. 2-5 μg antibody per milligrams of protein was added to the pre-cleared lysates and incubated on a rotator at 4°C overnight. Following over-night incubation Protein G Dyna beads were then added for 1h. Beads were washed four times in IP buffer, containing 300 mM NaCl and then resuspended in 40 μL of 2x Laemmli Sample Buffer and then boiled for 10min. Samples were then analyzed by SDS-PAGE and immunoblotting as described above. Antibodies used in the study is provided in the Supplementary table 3.

**Gene overexpression studies**

Plasmids expressing TMPRSS2 (pCSDest-hTMPRSS2, 53887) and SARS-CoV-2 spike protein (pLVX-EF1alpha-nCoV2019-S-2xStrep-IRES-Puro) were transfected alone or co-transfected into HEK-293T cells using Lipofectamine 2000 (Invitrogen Life Technologies, 12566014) as per manufacturer's protocol. 48-72h post-transfection the cells were harvested and subjected to immunoblotting. See Supplementary table 2 for the information on plasmid constructs used in the study.

**Spinoculation**

To evaluate the activity of TMPRSS2 on cell surface-bound SARS-CoV-2 Spike priming, the HIV/NanoLuc SARS-CoV-2 pseudo viral particles were spinoculated to HEK -293T-ACE2cl.22 cells. Prior to this, the cells were seeded in 6 well plates at a 70% confluency and transfected with pCSDest-hTMPRSS2 plasmid for 48h followed by 5h treatment with dmso or Camostat. For spinoculation, the cells were centrifuged in Sorvall Legend XTR Centrifuge (Thermo Scientific) for 1h at 800 x g, 37°C. Next, the cells were washed 2-3 times with the desired medium to remove the residual viral particles and incubated at 37°C and 5% Co2 for 4 more hours. Similarly, to check the effect of the endogenous TMPRSS2 on SARS-CoV-2 spike protein in LNCaP and Calu-3 cell lines, the same protocol was followed except the pCSDest-hTMPRSS2 transfection. After 4h incubation, the protein lysates were prepared, and the cleavage of SARS-CoV-2 Spike protein was analyzed through immunoblotting.

**Pseudo virus packaging**

To generate HIV/NanoLuc SARS-CoV-2 pseudo viral particles, HEK-293T cells were plated in 10 cm dish in 10ml DMEM + 10% FBS on day 0. Sixteen hours later, cells were starved with 4 ml of serum-free Opti-MEM (Thermo Fisher, 11058021) for 1 h. 6 μg of pHIV-1NL4-3 ΔEnv-NanoLuc reporter virus plasmid and 3 μg of SARS-CoV-2 spike protein plasmid (pSARS-CoV-2-SΔ19 or pLVX-EF1alpha-nCoV2019-S-2x Strep-IRES-Puro) were mixed thoroughly with 500 μl Opti-MEM. 9 μL of Lipofectamine 2000 (Thermo Fisher, 11668019) was diluted in 500 μl Opti-

MEM and mixed thoroughly. The plasmid and lipofectamine mixture were incubated together at room temperature for 15min before adding to the cells dropwise. After 8h of incubation, the media was replaced with 10ml of DMEM + 10% of FBS. After 48h and 60h post-transfection, the supernatant was harvested and clarified by centrifugation at 500 x g for 10min and filtered through a 0.45 µm syringe filter (VWR, 28145-481). To concentrate the virus, 1 volume of Lenti-X Concentrator (Takara, 631232) was mixed with 3 volumes of clarified supernatant and incubated at 4°C for 5 h, followed by centrifugation at 1,500 x g for 45min at 4°C. The viral pellet was resuspended in 1/5 of its original volume and stored at –80°C in single-use aliquots. HIV/NanoLuc VSV-G control viral particles were generated by merely replacing Spike S plasmid with VSV-G.

**Transduction and Nano-Luciferase Reporter Assay**

For SARS-CoV-2 Spike pseudovirus cell entry, 20,000 cells were seeded in 96-well plates on day 0 to achieve 50%-60% of confluency. On day 1, cells were treated with ARD-69 (500 nM), Enzalutamide (10 µM) or camostat mesylate (300 µM), respectively or camostat mesylate in combinations of ARD-69 or Enzalutamide in 50 µL of full media for one hour before transduction. To test effect the of androgen deprivation on transduction efficiency, 20,000 cells were seeded in 96-well plates on day 0 to achieve 50%-60% of confluency. On day 1, we genteelly washed the cells with PBS and changed the media to RPMI + 10% CSS. On day 3, we pretreated the cells with DHT (10 nM) and/or camostat mesylate (300 µM) for one hour before transduction.

For transduction, 50 µL of HIV/NanoLuc VSVG or HIV/NanoLuc SARS-CoV-2 pseudo viral particles were added to each well of the cells. The transduction efficiency (pseudovirus entry) was quantified 24 h post-transduction using Nano-Glo Luciferase Assay System (Promega, N1110). Cells were lysed in 50 µL of buffer for 5 min at room temperature, and then lysates were transferred to white opaque flat-bottom 96-well plates for luminescence reading on BioTek Synergy HT. Relative luminescence units (RLU) obtained were normalized to the values derived from cells infected with the SARS-CoV-2 pseudovirus with DMSO or DHT+DMSO treatment to present the relative entry efficiency.

**Mass Spectrometry Analysis**

Streptavidin IP eluate was resolved with SDS-PAGE and visualized with Coomassie stain. Regions corresponding to two prominent bands were excised from the gel and cut into 1 $mm^3$ cubes. They were destained with 50% Methanol/1.25% Acetic Acid, reduced with 5 mM DTT (Dithiothreitol) (Thermo), and alkylated with 40 mM iodoacetamide (Sigma). Gel pieces were then washed with 20 mM ammonium bicarbonate (Sigma) and dehydrated with acetonitrile (Fisher). Trypsin (Promega) (5 ng/mL in 20 mM ammonium bicarbonate) was added to the gel pieces and proteolysis was allowed to proceed overnight at 37ºC. Peptides were extracted with 0.3% triflouroacetic acid (J.T.Baker), followed by 50% acetonitrile. Extracts were combined and the volume was reduced by vacuum centrifugation.

Samples were analyzed on a QExactive HF mass spectrometer (Thermofisher Scientific San Jose, CA) coupled with an Ultimate 3000 nano UPLC system and EasySpray source. Peptides were separated by reverse phase (RP)-HPLC on Easy-Spray RSLC C18 2 um 75 µm id × 50cm column at 50ºC. Mobile phase A consisted of 0.1% formic acid and mobile phase B of 0.1% formic acid/acetonitrile. Peptides were eluted into the mass spectrometer at 300 nL/min with each RP-LC run comprising a 95-minute gradient from 1 to 3% B in 5 min, 3-45%B in 90 min. The mass spectrometer was set to repetitively scan m/z from 300 to 1800 (R = 120,000) followed by data-dependent MS/MS scans on the twenty most abundant ions, minimum AGC 1e4, dynamic exclusion with a repeat count of 1, repeat duration of 15s, (R=45000) and a NCE of 27. FTMS full scan AGC target value was 5e5, while MSn AGC was 1e5, respectively. MSn injection time was 120 ms; microscans were set at one. Rejection of unassigned, 1, 6-8 and >8 charge states was set.

**QA/QC and system suitability**

For online monitoring of the QExactive HF instrument, parallel reaction monitoring (PRM) analysis for the spiked-in iRT peptides was performed though Skyline AutoQC, and the data were uploaded and accessed in Skyline Panorama (Bereman et al., 2016). Meanwhile, as a measure for QC/QA, we injected standard E. coli protein digest in between samples (one injection after every four injections). The collected DDA data were analyzed in MaxQuant. The MaxQuant output was subsequently visualized using the PTXQC package to track the quality of the instrumentation(Bielow et al., 2016).

**Search MS raw files**

Protein and peptide identification/quantification was performed with MaxQuant (1.6.14.0) using a SARS COVID19 reference database from Uniprot (reviewed canonical and isoforms; downloaded on 20200904). Carbamidomethyl of Cys was defined as a fixed modification. Oxidation of Met and Acetylation of protein N-terminal were set as variable modifications. Trypsin/P was selected as the digestion enzyme, and a maximum of 3 labeled amino acids and 2 missed cleavages per peptide were allowed. Fragment ion tolerance was set to 0.5 Da. The MS/MS tolerance was set at 20 ppm. The minimum peptide length was set at 7 amino acids. The rest of the parameters were kept as default. The MaxQuant results were imported into Scaffold (Proteome Software) for data visualization.

**Table S1. List of Oligonucleotide Primers used in this study**

| SYBR Green qPCR | Sequence |
|---|---|
| mouse GAPDH FWD | CGACTTCAACAGCAACTCCCACTCTTCC |
| mouse GAPDH REV | TGGGTGGTCCAGGGTTTCTTACTCCTT |
| mouse AR FWD | GATGGTATTTGCCATGGGTTG |
| mouse AR REV | GGCTGTACATCCGAGACTTGTG |
| mouse TMPRSS2 FWD | GGCCGCTGGTTACTTTGAAG |
| mouse TMPRSS2 REV | TCGTGTCCCCAATCAGCC |
| mouse ACEII FWD | GGATACCTACCCTTCCTACATCAGC |
| mouse ACEII REV | CTACCCCACATATCACCAAGCA |
| human GAPDH FWD | TGCACCACCAACTGCTTAGC |
| human GAPDH REV | GGCATGGACTGTGGTCATGAG |
| human AR FWD | CAGTGGATGGGCTGAAAAAT |
| human AR REV | GGAGCTTGGTGAGCTGGTAG |
| human TMPRSS2 FWD | CAGGAGTGTACGGGAATGTGATGGT |
| human TMPRSS2 REV | GATTAGCCGTCTGCCCTCATTTGT |
| human ACEII FWD | CATTGGAGCAAGTGTTGGATCTT |
| human ACEII REV | GAGCTAATGCATGCCATTCTCA |

**Table S2. List of Plasmids used in this study**

| Plasmid | Source | Cat. # |
|---|---|---|
| pCSDest-hTMPRSS2 | Addgene | 53887 |
| pLVX-EF1alpha-nCoV2019-S-2xStrep-IRES-Puro | Gordon DE et al. Nature 2020 | - |
| pHIV-1(NL4.3) Env-NanoLuc | Gift from Paul D. Bieniasz, Rockefeller University | - |
| pSARS-CoV-2-S(D19) | Gift from Paul D. Bieniasz, Rockefeller University | - |
| VSV-G | Addgene | - |

**Table S3. List of Antibodies used in this study**

| Antibody | Application | Source | Cat. # |
|---|---|---|---|
| Mouse ACE2 | IHC/IB | R&D System | AF3437 |
| Human ACE2 | IB | R&D System | AF933 |
| TMPRSS2 | IB | Abcam | ab92323 |
| TMPRSS2 | IHC/IB | Abcam | ab109131 |
| AR | IHC/IB | Abcam | ab74272 |
| pS81-AR | IB | Millipore | 07-1375-EMD |
| HIV-1 core antigen-RD1, KC57 (p24) | IB | Beckman Coulter Life Sciences | 6604667 |
| Streptavidin-tag II | IB | Abcam | ab76950 |
| GAPDH | IB | Cell Signaling Technology (CST) | 3683S |
| β-Actin (D6A8) | IB | CST | 8457S |

Bereman, M.S., Beri, J., Sharma, V., Nathe, C., Eckels, J., MacLean, B., and MacCoss, M.J. (2016). An Automated Pipeline to Monitor System Performance in Liquid Chromatography-Tandem Mass Spectrometry Proteomic Experiments. J Proteome Res *15*, 4763-4769.

Bielow, C., Mastrobuoni, G., and Kempa, S. (2016). Proteomics Quality Control: Quality Control Software for MaxQuant Results. J Proteome Res *15*, 777-787.

Consortium, G.T., Laboratory, D.A., Coordinating Center -Analysis Working, G., Statistical Methods groups-Analysis Working, G., Enhancing, G.g., Fund, N.I.H.C., Nih/Nci, Nih/Nhgri, Nih/Nimh, Nih/Nida*, et al.* (2017). Genetic effects on gene expression across human tissues. Nature *550*, 204-213.

Heinz, S., Benner, C., Spann, N., Bertolino, E., Lin, Y.C., Laslo, P., Cheng, J.X., Murre, C., Singh, H., and Glass, C.K. (2010). Simple combinations of lineage-determining transcription factors prime cis-regulatory elements required for macrophage and B cell identities. Mol Cell *38*, 576-589.

Zhang, Y., Liu, T., Meyer, C.A., Eeckhoute, J., Johnson, D.S., Bernstein, B.E., Nusbaum, C., Myers, R.M., Brown, M., Li, W.*, et al.* (2008). Model-based analysis of ChIP-Seq (MACS). Genome Biol *9*, R137.
